# Transcription Factor Acj6 Controls Dendrite Targeting via Combinatorial Cell-Surface Codes

**DOI:** 10.1101/2021.11.03.467130

**Authors:** Qijing Xie, Jiefu Li, Hongjie Li, Namrata D Udeshi, Tanya Svinkina, Daniel Orlin, Sayeh Kohani, Ricardo Guajardo, DR Mani, Chuanyun Xu, Tongchao Li, Shuo Han, Wei Wei, S Andrew Shuster, David J Luginbuhl, Stephen R. Quake, Swetha E. Murthy, Alice Y Ting, Steven A Carr, Liqun Luo

## Abstract

Transcription factors specify the fate and connectivity of developing neurons. We investigate how a lineage-specific transcription factor, Acj6, controls the precise dendrite targeting of *Drosophila* olfactory projection neurons (PNs) by regulating the expression of cell-surface proteins. Quantitative cell-surface proteomic profiling of wild-type and *acj6* mutant PNs in intact developing brains and a proteome-informed genetic screen identified PN surface proteins that execute Acj6-regulated wiring decisions. These include canonical cell adhesion molecules and proteins previously not associated with wiring, such as Piezo, whose mechanosensitive ion channel activity is dispensable for its function in PN dendrite targeting. Comprehensive genetic analyses revealed that Acj6 employs unique sets of cell-surface proteins in different PN types for dendrite targeting. Combinatorial expression of Acj6 wiring executors rescued *acj6* mutant phenotypes with higher efficacy and breadth than expression of individual executors. Thus, Acj6 controls wiring specificity of different neuron types by specifying distinct combinatorial expression of cell-surface executors.

## INTRODUCTION

The nervous system is incredibly complex. An adult human brain consists of an average of about 86 billion neurons (Azevedo et al., 2009) forming ∼10^14^ connections. Even in simpler organisms such as the fruit fly, there are around 200,000 neurons in its brain (Raji and Potter, 2021) forming ∼10^7^ connections. Precise formation of these neuronal connections ensures proper information processing, which underlies all nervous system functions. Decades of research have identified numerous wiring molecules that specify these neuronal connections.

Most molecules that instruct neuronal wiring discovered thus far fall into two categories— transcription factors and cell-surface proteins (Butler and Tear, 2007; Jan and Jan, 2010; Kolodkin and Tessier-Lavigne, 2011; Sanes and Zipursky, 2020; Santiago and Bashaw, 2014). Transcription factors are central commanders specifying cell fate, morphology, and physiology while cell-surface proteins execute these commands through interaction with cellular environment. In developing neurons, it is presumed that transcription factors control wiring specificity through regulation of cell-surface protein expression. However, causal links between transcription factor and cell-surface wiring proteins have only been established in a few isolated cases (Butler and Tear, 2007; Lee et al., 2015; Peng et al., 2018; Peng et al., 2017; Santiago and Bashaw, 2014; 2017). Moreover, these reports all focus on the regulation of one or two known wiring regulators in a specific neuron type. Therefore, the number and identity of cell-surface protein(s) a transcription factor regulates to execute its wiring command remain largely unknown (Figure 1A, top). It is also unclear whether a transcription factor regulates the same set or different sets of cell-surface proteins in different neuron types to specify their connectivity (Figure 1A, bottom).

**Figure 1.**
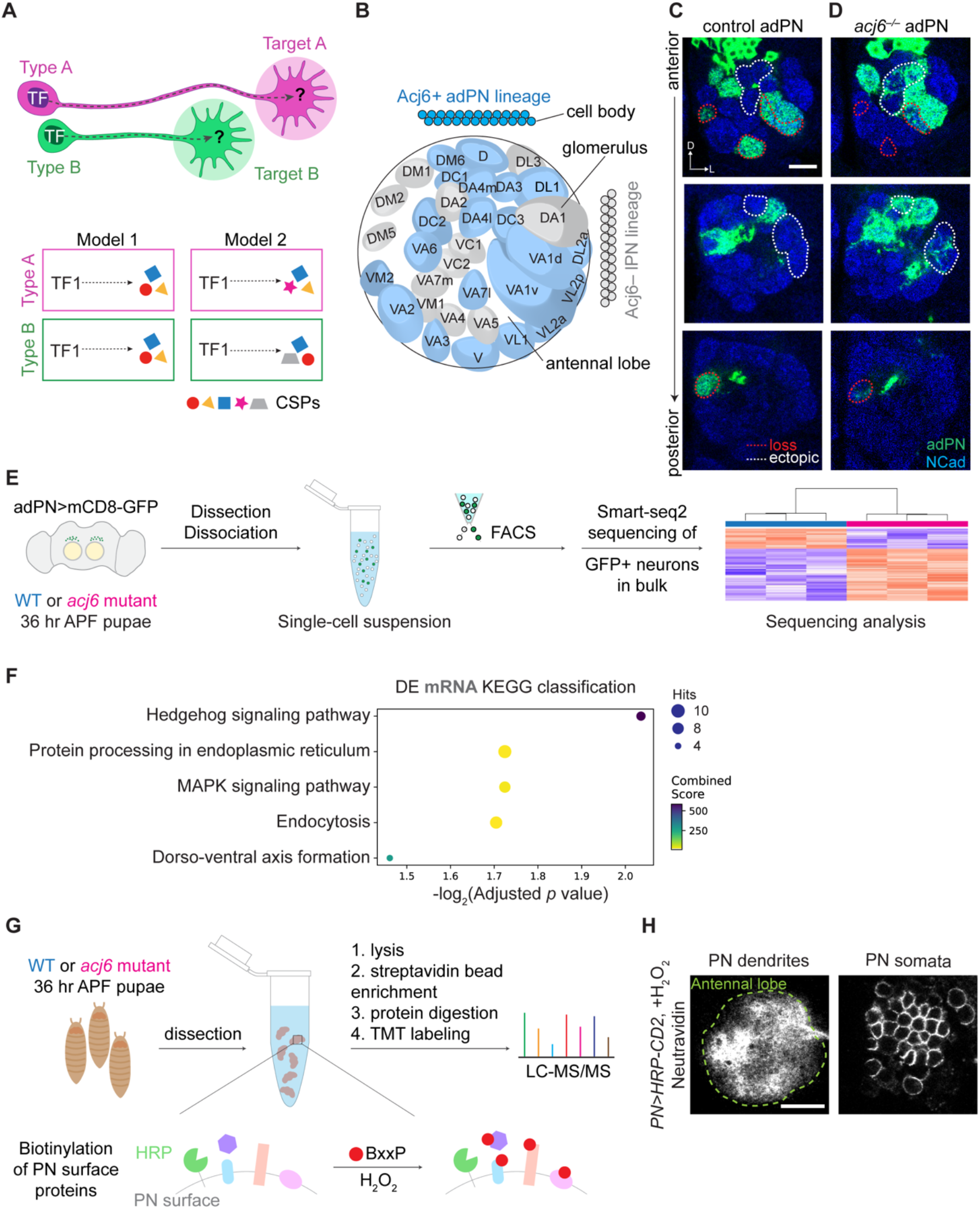
Identification of Acj6-regulated mRNAs and cell-surface proteins. (A) Central question: how does a transcription factor (TF) control targeting specificity of different neuron types by regulating the expression of cell-surface proteins (CSPs)? (B) Diagram of cell body locations and glomerular innervation patterns of *Drosophila* olfactory projection neurons (PNs) derived from either the anterodorsal (adPN, colored in blue) or the lateral (lPN) neuroblast lineage. Acj6 is specifically expressed in adPNs, which target dendrites to a stereotyped subset of glomeruli (colored in blue). (C and D) Compared to those of wild-type (C), dendrites of *acj6^−/−^* adPN neuroblast clones (D) exhibit both loss of innervation (circled in red) and ectopic targeting (circled in white). Blue, anti-NCad antibody staining revealing glomerular organization of the antennal lobe. Green, anti-GFP staining of membrane-targeted GFP expressed by PNs. (E) Schematic of adPN transcriptomic profiling in developing wild-type and *acj6* mutant brains. APF, after puparium formation. (F) Top five KEGG pathways enriched among genes that are differentially expressed (DE; adjusted *p* < 0.05) between wild-type and *acj6* mutant adPNs. (G) Schematic of PN surface proteomic profiling in developing wild-type and *acj6* mutant brains. (H) Neutravidin staining of developing PNs expressing HRP after the cell surface biotinylation reaction. Signals are absent intracellularly in PN somata. Scale bars, 20 μm. D, dorsal; L, lateral. See also Figure S1.

Here, we systematically addressed these questions using the transcription factor Acj6 as an example. Acj6 is a member of POU domain transcription factors, which are widely used from *C. elegans* to mammals to regulate neural development (Certel et al., 2000; Komiyama et al., 2004; Komiyama et al., 2003; Schonemann et al., 1998). In the *Drosophila* olfactory system, about 50 types of cholinergic excitatory PNs are derived from two distinct neuroblast lineages (Jefferis et al., 2001). Each PN type targets dendrites to a stereotyped antennal lobe glomerulus according to its lineage and birth order (Jefferis *et al*., 2001; Lin et al., 2012; Marin et al., 2005; Yu et al., 2010). Acj6 is specifically expressed in postmitotic neurons in the anterodorsal PN (adPN) lineage (Figure 1B), whereas another POU domain transcription factor Vvl is expressed in the lateral PN (lPN) lineage, to instruct lineage-specific dendrite targeting of adPNs and lPNs (Komiyama *et al*., 2003).

In this study, we profiled the developing PN-surface proteomes of wild-type and *acj6*- mutant brains by quantitative liquid chromatography-tandem mass spectrometry (LC-MS/MS) and performed a proteome-informed genetic screen to identify cell-surface proteins that execute Acj6’s wiring commands. We discovered that Acj6 instructs PN dendrite targeting by regulating the expression of both canonical wiring molecules with extracellular cell-adhesion domains and unconventional molecules that have been studied mostly in the context of neuronal function, such as the mechanosensitive ion channel Piezo. Overexpressing one or a pair of Acj6 target(s) downregulated on the *acj6* mutant PN surface gave rise to specific and combinatorial rescue of *acj6* mutant phenotypes, indicating that Acj6 regulates unique combinations of cell-surface proteins in different PN types to specify distinct dendrite targeting. Together, we provide the most comprehensive evidence to date of how a single transcription factor can specify many unique neuronal connections and present functional data showing that neuronal wiring is controlled by a combinatorial code of cell-surface molecules.

## RESULTS

### Acj6 shapes the PN surface proteomic milieu

Acj6 is required for the proper dendrite targeting of adPNs, as *acj6* mutant adPNs exhibit both loss of innervation in glomeruli normally targeted by adPN dendrites and ectopic innervation in glomeruli normally not targeted by these adPN dendrites (Komiyama *et al*., 2003; Figures 1C and 1D). Acj6 may regulate dendrite targeting through direct transcriptional regulation of cell-surface proteins or through intermediate transcription factors and post-transcriptional mechanisms. To investigate this, we performed transcriptome profiling of wild-type and *acj6* mutant adPNs at 36– 40h after puparium formation (APF), when PN dendrites are actively making wiring decision (Figure 1E; Table S1). We found that genes involved in protein processing in the endoplasmic reticulum and endocytosis were among the top five pathways regulated by Acj6 (Figure 1F), suggesting that Acj6 could shape the PN-surface proteomic landscape post-transcriptionally (Figure S1).

Therefore, to capture the entirety of Acj6-regulated cell-surface proteins, we performed PN-surface proteomic profiling in intact wild-type or *acj6* mutant fly brains (Figure 1G) at 36–40h APF utilizing our recently developed in situ cell-surface proteomic profiling method (Li et al., 2020). Briefly, membrane-tethered horseradish peroxidases (HRP) expressed on PNs convert the membrane-impermeable biotin-xx-phenol (BxxP) substrate to phenoxyl radicals in the presence of H_2_O_2_ (Loh et al., 2016), which promiscuously biotinylate proteins on the PN surface. This approach led to biotinylation of PN-surface proteins with high spatial specificity (Figure 1H).

We applied an 8-plex tandem mass tag (TMT)-based quantitative mass spectrometry strategy to identify those biotinylated proteins (Li et al., 2020; Thompson et al., 2003) (Figure 2A). For each genotype, in addition to two biological replicates, we also included two negative controls, in which either H_2_O_2_ or HRP was omitted to account for non-specific bead binders, endogenously biotinylated proteins, and labeling by endogenous peroxidases. Each sample (derived from ∼1100 dissected pupal brains) was separately lysed and enriched with streptavidin beads (Figure 2B). After on-bead trypsin digestion and TMT labeling, all samples were pooled for mass spectrometry analysis (Table S2).

**Figure 2.**
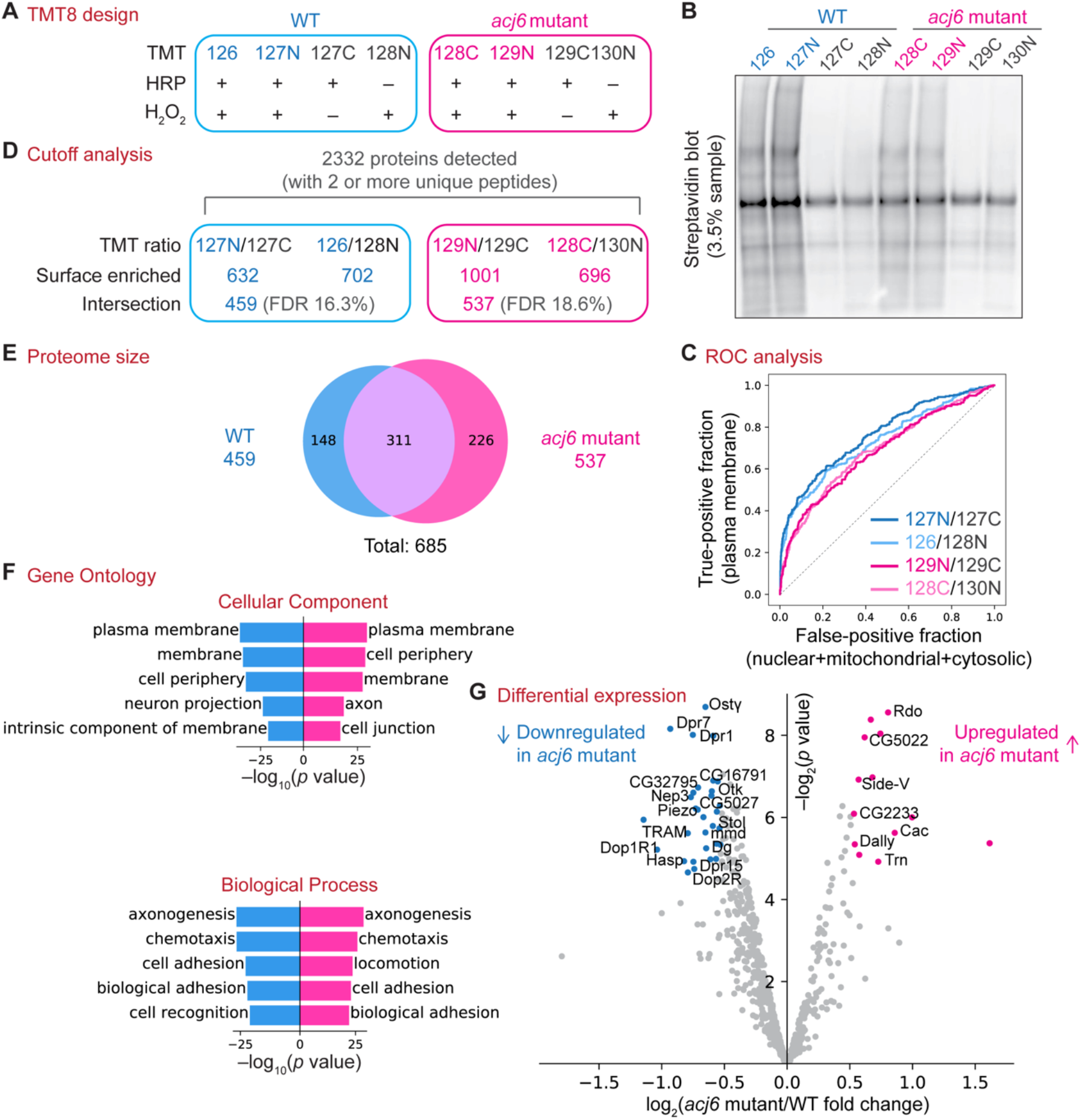
Cell-surface proteomes of wild-type and *acj6* mutant PNs. (A) Design of the 8-plex tandem mass tag (TMT)-based quantitative proteomic experiment. Each genotype comprises two biological replicates (blue or pink) and two negative controls omitting either HRP or H_2_O_2_ (black). Labels in the TMT row indicate the TMT tag used for each condition. (B) Streptavidin blot of the post-enrichment bead eluate—3.5% from each sample listed in (A). (C) Receiver operating characteristic (ROC) analysis showing the separation of true positives from false positives in each biological replicate using the experimental-to-control TMT ratio. (D) Summary of the cutoff analysis. Proteins with TMT ratios greater than the cutoffs in both biological replicates were retained (‘Intersection’). FDR: false discovery rate. (E) Venn diagram showing the size of and overlap between wild-type and *acj6* mutant PN surface proteomes. (F) Top five Cellular Component and Biological Process Gene Ontology terms for wild-type (blue) and *acj6* mutant (pink) PN surface proteomes. (G) Volcano plot showing proteins with altered protein levels on *acj6* mutant PN surface compared to wild-type PN surface. Proteins downregulated in *acj6* mutant [log_2_(fold change) less than –0.53 and *p*-value less than 0.05] are colored in blue. Proteins upregulated in *acj6* mutant [log_2_(fold change) greater than 0.53 and *p*-value less than 0.05] are colored in red. See also Figures S1 and S2.

Biological replicates in both genotypes showed high correlation (Figure S2A), illustrating the robustness of our method. Additionally, loss of Acj6 did not cause global disruption of protein expression (Figure S2B). We ranked proteins by their experimental-to-control TMT ratios in a descending order and found that known plasma membrane proteins were enriched while contaminants were sparse at the top of those ranked lists (bottom left corner, Figure 2C). Therefore, we filtered out contaminants using experimental-to-control TMT ratios (Hung et al., 2014; Li *et al*., 2020) (Figure 2D; Figure S2C), yielding PN surface proteomes of 459 and 537 proteins for wild-type and *acj6* mutant, respectively (Figure 2E). Cellular Component terms classified by Gene Ontology analysis showed that both proteomes consisted of proteins localized at the cell surface, confirming the spatial specificity of our approach (Figure 2F, top). Top Biological Process terms showed enrichment of neural developmental processes in both proteomes (Figure 2F, bottom), matching the developmental stage at which we conducted PN surface profiling.

Many proteins exhibited altered expression on the *acj6* mutant PN surface (Figure 2G). These include proteins belonging to classic cell adhesion protein families—Off-track (Otk), Sidestep-V (Side-V), and Dprs in the immunoglobulin (Ig) superfamily, and Reduced ocelli (Rdo) and Tartan (Trn) in the leucine-rich repeat superfamily. We also observed several proteins known to participate in neuronal function—the mechanosensitive ion channel Piezo, the dopamine receptors Dop1R1 and Dop2R, a regulator of voltage-gated calcium channel Stolid (Stol), and the voltage-gated calcium channel subunit α_1_ Cacophony (Cac). Most of these proteins were also identified using a more stringent cutoff criterion (Figures S2D–S2F). Consistent with our RNA-seq analysis suggesting post-transcriptional regulation of cell-surface proteins by Acj6 (Figure 1F), Acj6 binding site prediction suggested that most genes encoding cell-surface proteins differentially expressed in wild-type and *acj6* mutant were not direct targets of Acj6 (Figure S1D).

### A proteome-informed genetic screen identified new wiring molecules for PN dendrites

Among the Acj6-regulated PN surface proteins we identified, Trn is the only one known to participate in PN dendrite targeting (Hong et al., 2009). Therefore, we designed a genetic screen to systematically identify Acj6 targets with wiring functions.

We used a split GAL4 intersectional strategy (Luan et al., 2006; Pfeiffer et al., 2010) to reconstitute full-length GAL4 in around half of all adPN types by combining the PN-specific *VT033006-GAL4^DBD^* with the adPN lineage-restricted *C15-p65^AD^*. This “*adPN-GAL4*” line was used to knock down (with *UAS-RNAi*) or overexpress candidates (with *UAS-cDNA*) whose expressions were down- or up-regulated on the *acj6* mutant PN surface, respectively. Due to the limited availability of existing *UAS-cDNA* lines for overexpression, most candidates we tested were downregulated in *acj6* mutant, which means that Acj6 normally promotes their expression.

We scored the innervation degrees of 38 glomeruli in each antennal lobe of each genotype (Figure 3A; Figure S3). To determine whether altering the expression of these Acj6-regulated PN surface proteins caused significant dendrite targeting changes, the innervation extent of each glomerulus in each genotype was compared to control using a Chi-squared test (Figure 3A; Figure S4). In contrast to our previous wiring molecule screening schemes (Ward et al., 2015; Xie et al., 2019), this screening and scoring strategy can monitor dendrite targeting defects in many PN types simultaneously instead of just two to three PN types located in a specific region. This is critical as Acj6 controls dendrite targeting of many PN types and might regulate expression of different wiring molecules in different PN types.

**Figure 3.**
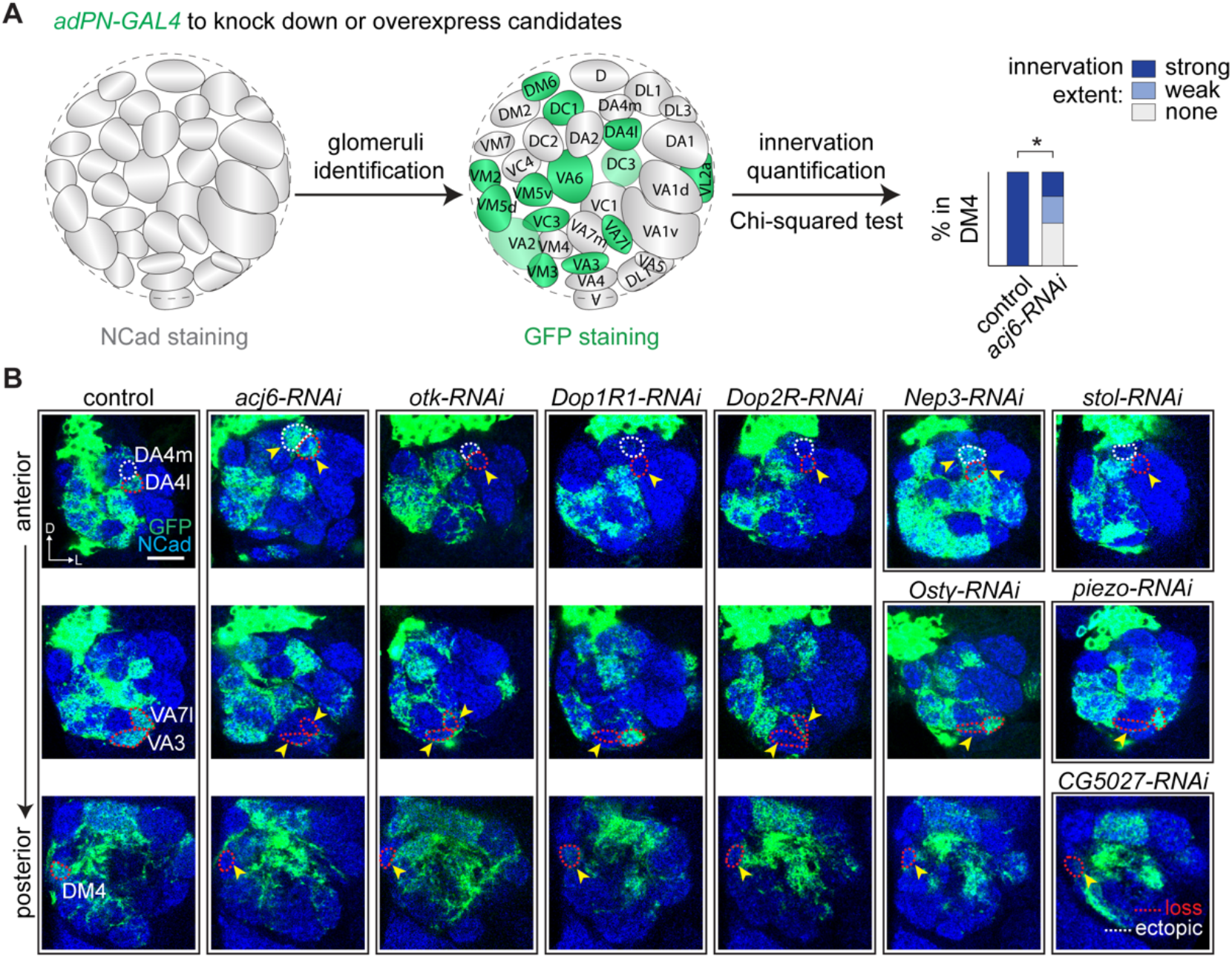
A proteome-informed genetic screen to identify wiring executors of Acj6. (A) Schematic of the genetic screen and quantification. Dendrite innervation of each glomerulus was scored into three categories: strong, weak, and none. Scorer was blind to genotypes. (B) Confocal images showing dendrite innervation patterns of *adPN-GAL4+* PNs (green) in controls and when *acj6* or candidate genes were knocked down by RNAi. Blue, N-cadherin (NCad) staining highlight the glomeruli. Glomeruli with decreased PN dendrite innervation are circled in red while ectopically targeted glomeruli are circled in white. Arrowheads point to the glomeruli exhibiting mistargeting phenotypes. Scale bars, 20 μm. D, dorsal; L, lateral. See also Figures S3 and S4.

We identified many new molecules required for PN dendrite targeting (Figure 3B). These include Otk, Neprilysin 3 (Nep3), and Dystroglycan (Dg), which have been previously shown to be required for neurite targeting of other neuron types (Cafferty et al., 2004; Li *et al*., 2020; Shcherbata et al., 2007; Winberg et al., 2001). In addition, knockdown of *Dop1R1*, *Dop2R, stol*, and *piezo,* which are traditionally thought to mediate neuronal function instead of development, also caused PN wiring defects. Previously uncharacterized genes, such as *CG5027*, also contributed to the targeting accuracy of PN dendrites.

As with knocking down *acj6* itself, knocking down Acj6-regulated cell-surface proteins caused abnormal targeting of adPN dendrites of many glomeruli. Notably, these ectopic targeting or loss of targeting resembled *acj6-RNAi* in multiple glomeruli including DA4m, DA4l, VA7l, VA3, and DM4 (Figure 3B). Furthermore, dendrite targeting to an individual glomerulus can be affected by knocking down several different molecules. For example, loss of VA7l innervation was observed when knocking down *acj6*, *otk*, or *Dop2R*, but not *piezo*, *Dop1R1*, or *ostγ* (second row of Figure 3B). These data suggest that many Acj6-regulated cell-surface proteins indeed regulate wiring specificity, and that proper targeting of some PN dendrites requires multiple cell-surface proteins controlled by Acj6. Additionally, knockdown of some candidates also caused unique dendrite mistargeting patterns not observed in *acj6-RNAi* (Figure S4), suggesting that these molecules also have wiring functions independent of Acj6.

### Ig-superfamily protein Otk is a cell-surface wiring executor for Acj6

To establish causal relationships between Acj6 and its targets in regulating wiring specificity (Figure 4A), we tested whether forced expression of specific Acj6-downregulated cell-surface proteins can rescue *acj6* mutant phenotypes using mosaic analysis with a repressible cell marker (Lee and Luo, 2001) (MARCM; Figures S5A and S5B) with null mutant alleles and full-length *UAS-cDNA* transgenes. Besides bypassing Acj6’s transcriptional regulation, the amplification effect of the GAL4/UAS system makes it likely that forced expression of candidate cell-surface proteins would also overcome the post-transcriptional regulation by Acj6.

**Figure 4.**
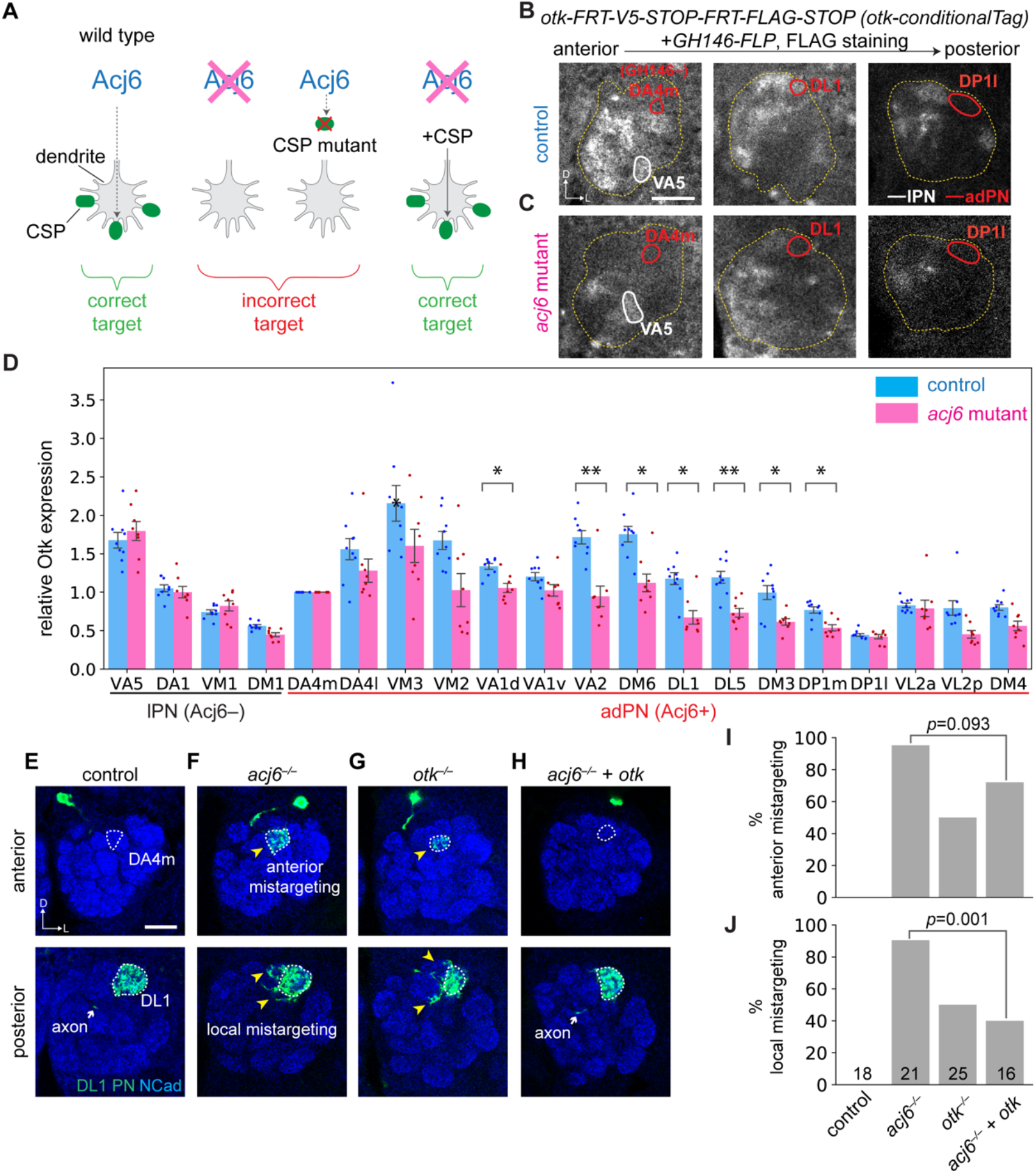
Off-track (Otk) is an Acj6 executor for instructing PN dendrite targeting. (A) Criteria of Acj6’s cell-surface wiring executors. Loss of either Acj6 or its cell-surface wiring executor would lead to similar dendrite mistargeting. Supplying back the cell-surface wiring executor of Acj6 would rescue *acj6* mutant phenotype. (B and C) Expression of Otk in PNs at 42–48 hr APF in wild-type (B) or *acj6* mutant (C) brains. Antennal lobe (outlined in yellow) and the DA4m (GH146– adPN), VA5 (lPN), and DL1 (adPN) glomeruli are outlined. *GH146-FLP* was used to express FLP in the majority of PNs. (D) Quantification of Otk expression in developing PNs of wild-type (n = 9) and *acj6-*mutant (n = 8) animals. Fluorescence intensities were normalized to the *GH146-FLP* negative DA4m glomerulus in each antennal lobe. Mean ± s.e.m. *: *p* < 0.05; **: *p* < 0.01 (two-tailed t-test; *p*- values were adjusted for multiple comparisons using Bonferroni’s method). Note that the fluorescence intensity is always lower in deeper (more posterior) sections due to tissue scattering, so the absolute intensity cannot be compared across different PN types. (E) Dendrites of wild-type DL1 single-cell clones innervate exclusively the posterior DL1 glomerulus (n = 18). (F) *acj6^−/−^* DL1 single-cell clones mistarget to both nearby glomeruli and a distant anterior glomerulus (n = 21). Yellow arrowhead, mistargeted PN dendrites. (G) *otk^−/−^* DL1 single-cell clones phenocopy *acj6^−/−^* (n = 25). (H) *acj6^−/−^* DL1 dendrite mistargeting phenotypes were partially rescued by overexpressing *otk* (n = 16). (I and J) Quantification of DL1 single-cell clones that mistargeted at a distance (I) and locally (J). *p*-values of Chi-squared tests are shown on plots. Scale bars, 20 μm. D, dorsal; L, lateral. Arrow, PN axons. See also Figures S5.

We first investigated Otk, a transmembrane protein containing five extracellular Ig-like domains implicated in axon guidance in embryonic motor axons (Winberg *et al*., 2001) and photoreceptor axons (Cafferty *et al*., 2004). We generated a conditional tag to examine endogenous Otk expression specifically in PNs (Figures S5C–S5E). We found that Otk was normally expressed in subsets of both Acj6-positive adPNs and Acj6-negative lPNs during development (Figure 4B). In *acj6* mutant, there was an apparent overall decrease in Otk signal (Figure 4C). Quantification of fluorescence intensity in individual glomeruli revealed that Otk expression was indeed decreased in many adPNs such as DL1, but not in lPNs such as VA5 (Figure 4D). However, some adPN types such as VA1v and VL2a had similar levels of Otk expression in *acj6* mutant and wild-type animals (Figure 4D), and some adPN types such as DP1l had barely detectable Otk expression in wild-type animals (Figure 4B). Thus, Acj6 promotes Otk expression on the surface of a subset of adPNs.

We performed MARCM analysis by deleting *otk* in either anterodorsal or lateral neuroblast clones (Figure S5B) and found dendrite mistargeting in both (Figures S5F–S5I), consistent with *otk* expression in both Acj6+ adPNs and Acj6– lPNs. These data validated Otk as a wiring molecule for PN dendrites. To test if Acj6 regulates Otk expression to instruct adPN dendrite targeting, we focused on single-cell clones of DL1 PNs (Figure S5B). In wild-type animals, dendrites of DL1 PNs are confined to a specific glomerulus located in a posterior section of the antennal lobe (Figure 4E, outlined in the bottom panel). Deleting *acj6* or *otk* in DL1 PNs yielded nearly identical mistargeting phenotypes: DL1 PN dendrites mistargeted to both nearby posterior glomeruli as well as the distant DA4m glomerulus in the anterior antennal lobe (Figures 4F and 4G). Furthermore, both local and long-range mistargeting phenotypes in *acj6* mutant were partially rescued by overexpressing *otk* in DL1 single cell clones (Figures 4H–4J). This incomplete rescue is consistent with the observation that *otk^−/−^* clones had lower phenotypic penetrance than *acj6^−/−^* clones (Figures 4I and 4J) and suggests that Acj6 controls the expression of additional wiring molecule(s) in DL1 PNs to ensure precise dendrite targeting.

These single-cell clone analyses, together with the observation that Otk expression was decreased in *acj6* mutant DL1 PNs (Figures 4B–4D), demonstrated that Acj6 directs precise targeting of DL1 PN dendrites in part by cell-autonomously promoting Otk expression.

### Piezo executes Acj6’s wiring command independently of its the mechanosensitive ion channel activity

An unexpected observation in our genetic screen was that knocking down the mechanosensitive ion channel, Piezo, disrupted normal PN dendrite targeting (Figure 3B; Figure S4). MARCM analysis using a *piezo* null mutant confirmed that both *acj6^−/−^* and *piezo^−/−^* adPNs in neuroblast clones mistargeted their dendrites to the DL2 and VL2a glomeruli (Figures 5A–5C), albeit with a lower penetrance in *piezo^−/−^* PNs (Figure 5I). *piezo* endogenous conditional tag revealed that Piezo was expressed in a sparse set of PNs, and expression of Piezo in some adPNs was downregulated in *acj6* mutant animals (Figures S6A and S6B). Furthermore, overexpressing wild-type *piezo* in adPN neuroblast clones not only rescued *piezo* mutant phenotypes (Figures 5E and 5I), but also partially rescued *acj6* mutant phenotypes (Figures 5D and 5I). Thus, Piezo is an executor for Acj6 in controlling dendrite targeting.

**Figure 5.**
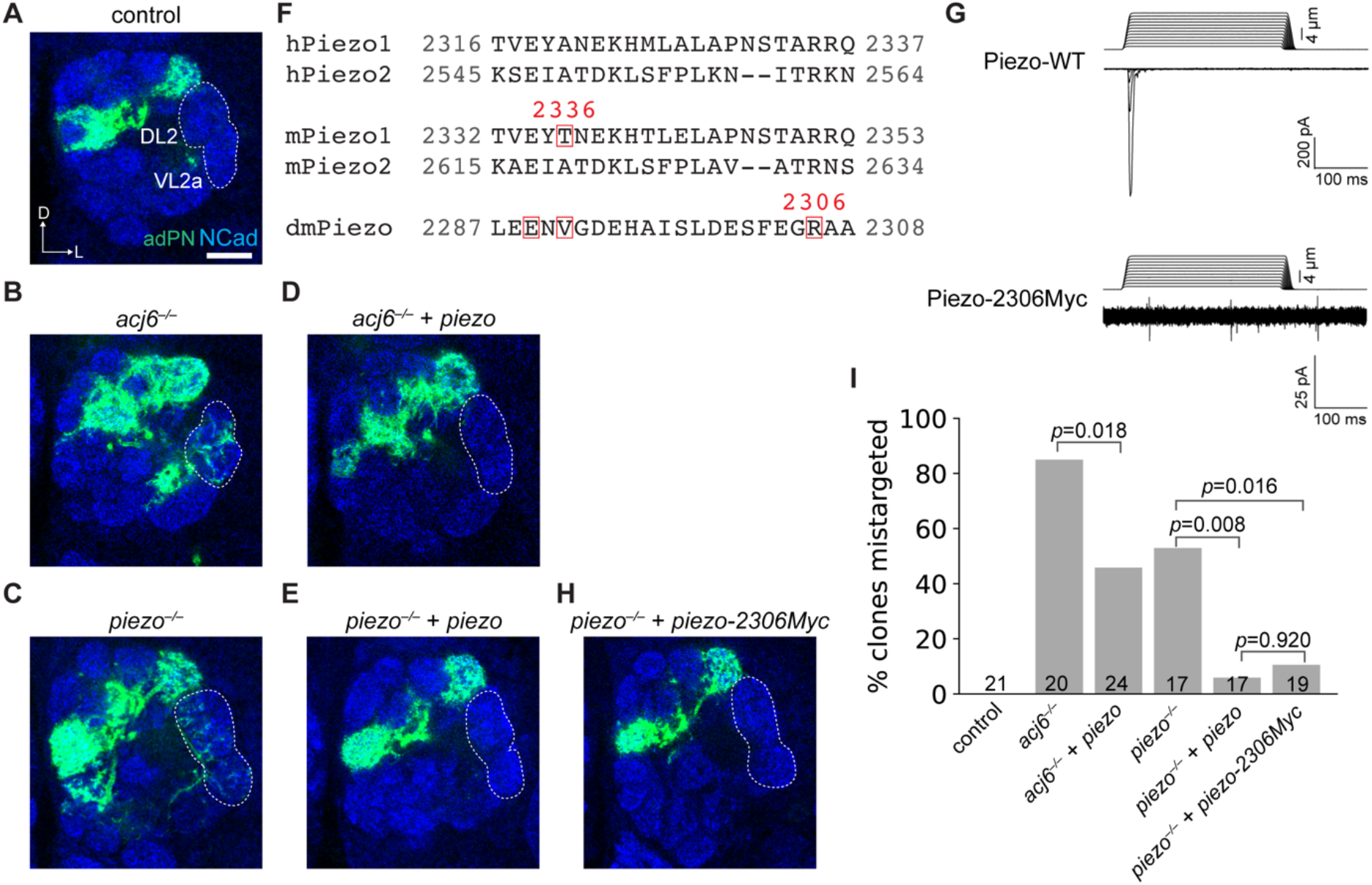
Piezo is an Acj6 executor for instructing PN dendrite targeting. (A–C) Dendrite innervation patterns of wild-type (n = 21) (A), *acj6^−/−^* (n = 20) (B), and *piezo^−/−^* (n = 17) (C) adPN neuroblast clones. DL2 and VL2a glomeruli are ectopically innervated in *acj6* and *piezo* mutants. (D and E) *acj6^−/−^* (D, n = 24) and *piezo^−/−^* (E, n = 17) mutant phenotypes were rescued by overexpressing wild-type *piezo*. (F) Sequence alignment of human Piezo1 (hPiezo1), human Piezo2 (hPiezo2), mouse Piezo1 (mPiezo1), mouse Piezo2 (mPiezo2), and fruit fly Piezo (dmPiezo). Red boxes highlight amino acids located N-terminal to Myc insertions. (G) A Myc tag inserted C-terminal to amino acid position 2306 of fly Piezo abolishes its mechanically activated channel activity. Representative traces of indentation-induced whole-cell currents from *piezo-WT* or *piezo-2306Myc* transfected human *PIEZO1* knockout HEK293T cells are shown (see Figure S6 for quantification as well as additional data on other Myc insertion mutants). (H) *piezo^−/−^* (n = 19) mutant phenotypes were rescued by overexpressing *piezo-2306Myc*. (I) Quantification of adPN dendrites in neuroblast clones that mistargeted to DL2 and VL2 glomeruli. *p*-values of Chi-squared tests are shown. Scale bars, 20 μm. D, dorsal; L, lateral. See also Figures S6.

Is the mechanosensitive ion channel activity of Piezo required for instructing PN dendrite targeting? A previous study showed that inserting a Myc tag at amino acid position 2336 in mouse Piezo1 abolishes its channel activity (Coste et al., 2015). Therefore, we inserted Myc tags at three amino acid locations in fly Piezo in a region homologous to the mouse Piezo1 2336 position (Figure 5F). All three Myc-insertion variants displayed proper cell-surface expression (Figure S6C). However, whole-cell patch clamp recordings showed that all three variants exhibited reduced mechanically activated currents in response to membrane indentation compared with control, and the mechanosensitive ion channel activity of Piezo-2306Myc was completely abolished (Figure 5G; Figures S6D and S6E).

Remarkably, overexpressing *piezo-2306Myc* in adPN neuroblast clones rescued dendrite targeting deficits of *piezo* mutants to the same extent as overexpressing wild-type *piezo* (Figures 5H and 5I). We further examined the Acj6-negative lPNs and observed that loss of Piezo also caused dendrite mistargeting, which was partially rescued by either wild-type Piezo or Piezo- 2306Myc (Figures S6F–S6I). This indistinguishable rescuing capacity suggested that Piezo regulates PN dendrite targeting independently of its mechanosensitive ion channel activity.

### Acj6 employs distinct cell-surface executors in different PN types

Data presented so far indicate that Acj6 directs the precise targeting of DL1 PN dendrites in part through Otk (Figure 4) and prevents mistargeting of adPN dendrites to the DL2 and VL2a glomeruli in part through Piezo (Figure 5). To investigate more cell-surface wiring executors of Acj6 in a broader set of PNs, we overexpressed in *acj6^−/−^* adPN neuroblast clones six additional cell-surface proteins that were downregulated in *acj6* mutant and examined dendrite innervation patterns in a broader set of PNs.

Overexpression of seven out of eight of these cell-surface proteins rescued specific subsets of *acj6* mutant phenotypes to different extents (Figure 6; Figure S7A), including ectopic targeting (DA4m, VA6, and DL2; Figures 6A–6F) and loss of innervation (VM2, VA1v, and VM7; Figures 6G–6L). The mistargeting of each PN type was rescued by specific cell-surface proteins. For example, loss of VA1v targeting in *acj6* adPN neuroblast clones was rescued by Nep3 (Figures 6I and 6J) while loss of VM7 targeting was rescued by the dopamine receptors Dop1R1 and Dop2R (Figures 6K and 6L). From the perspective of individual cell-surface protein, each participated in the targeting of specific glomeruli. For instance, Dg overexpression rescued mistargeting to VA6 and DL2 glomeruli (Figures 6C–6F), whereas Dop1R1 overexpression rescued VM7 targeting and mistargeting to DA4m in *acj6^−/−^* adPN neuroblast clones (Figures 6A, 6B, 6K, and 6L). Thus, Acj6-regulated cell-surface executors exhibited a complex but highly specific correspondence to individual adPN types and their target glomeruli in the developing antennal lobe.

**Figure 6.**
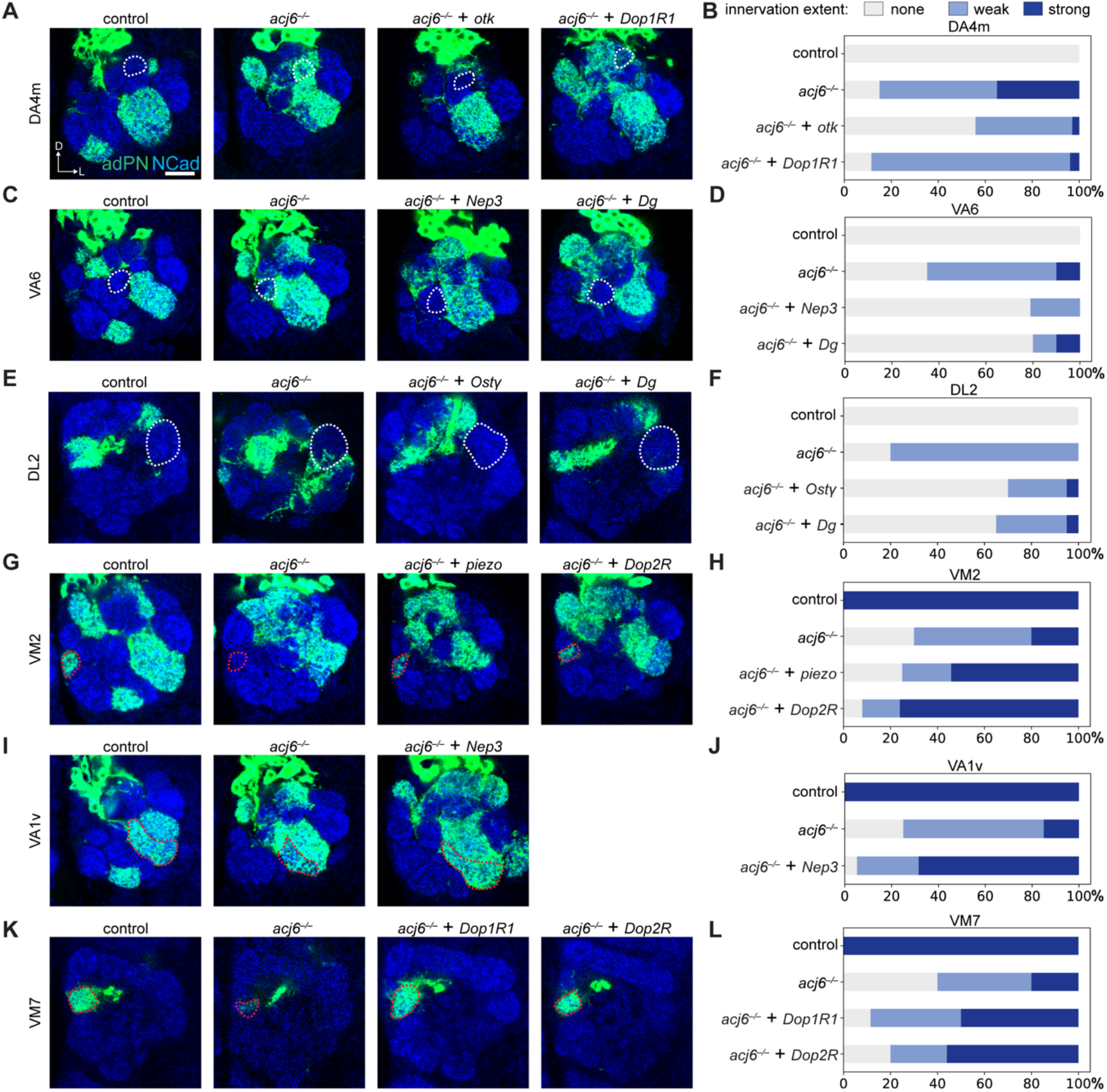
*acj6* mutant phenotypes rescued by individual cell-surface executors. (A and B) Ectopic targeting of adPN dendrites in neuroblast clones (adPN dendrites hereafter) to the DA4m glomerulus in *acj6* mutant was partially rescued by overexpressing *otk* or *Dop1R1*. (C and D) Ectopic targeting of adPN dendrites to the VA6 glomerulus in *acj6* mutant was partially rescued by overexpressing *Nep3* or *Dg*. (E and F) Ectopic targeting of adPN dendrites to the DL2 glomerulus in *acj6* mutant was partially rescued by overexpressing *Ostγ* or *Dg*. (G and H) Loss of adPN dendrite innervation in the VM2 glomerulus in *acj6* mutant was partially rescued by overexpressing *piezo or Dop2R*. (I and J) Loss of adPN dendrite innervation in the VA1v glomerulus in *acj6* mutant was partially rescued by overexpressing *Nep3*. (G and H) Loss of adPN dendrite innervation in the VM7 glomerulus in *acj6* mutant was partially rescued by overexpressing *Dop1R1 or Dop2R*. Bar graphs show fractions of clones belonging to three categories of innervation extent: strong, weak, and none. Scale bars, 20 μm. D, dorsal; L, lateral. See also Figures S7.

To quantitatively assess whether overexpression of Acj6-regulated cell-surface proteins indeed improved wiring precision of *acj6^−/−^* adPN dendrites, we performed Chi-squared tests comparing the innervation extent of each glomerulus in each rescue experiment to that in control or *acj6^−/−^* (Figure 7A). As shown in each row of Figure 7A, overexpressing a single Acj6-regulated cell-surface protein rescued a subset of *acj6^−/−^* phenotypes, consistent with our observation that loss of one cell-surface protein resembled only a subset of *acj6^−/−^* phenotypes (Figures 3–5).

**Figure 7.**
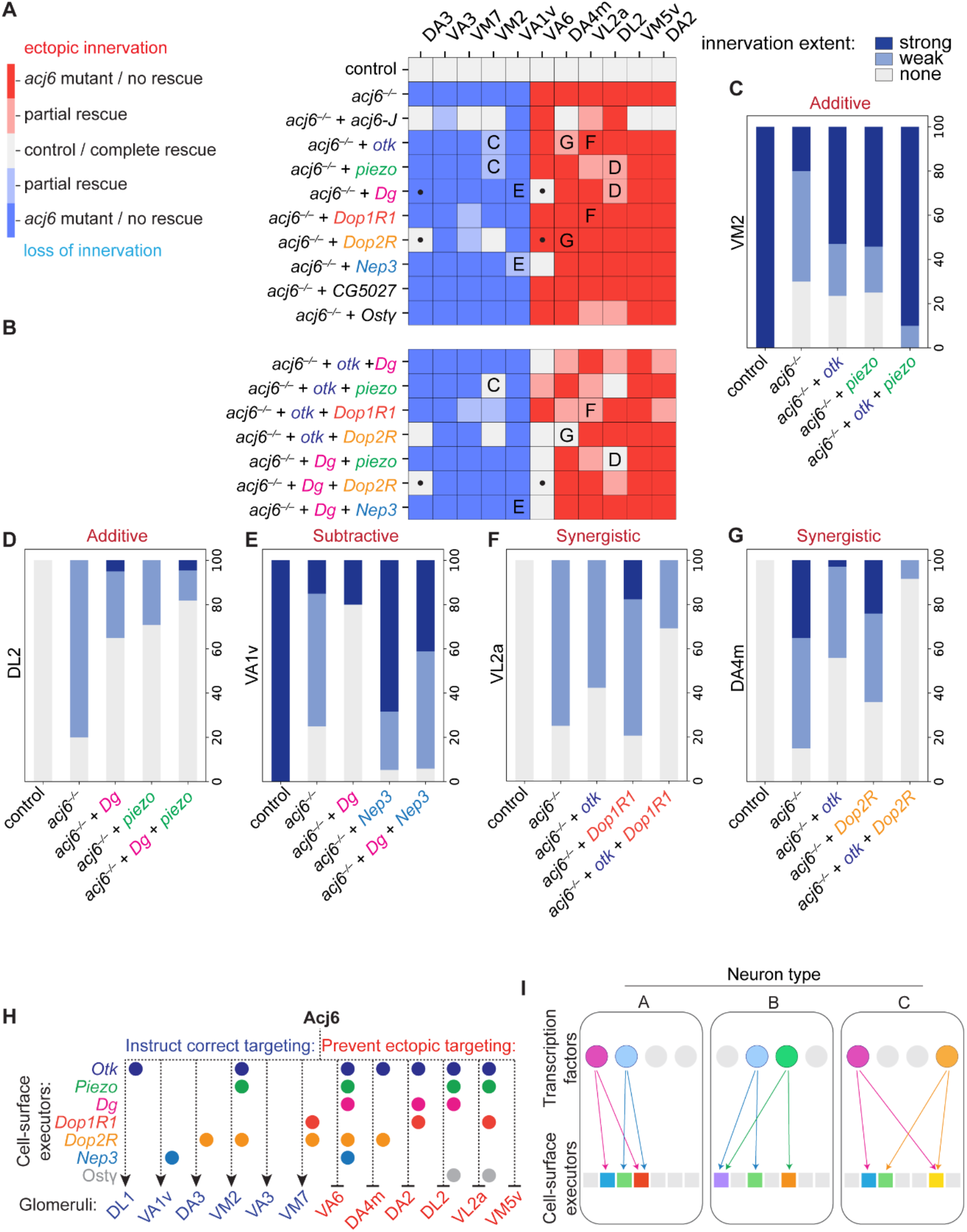
Acj6 instructs PN wiring through cell-surface combinatorial codes. (A and B) Heatmap summarizing changes in the dendrite innervation pattern of adPN neuroblast clones to eleven glomeruli (columns). Dark red (ectopic innervation) and dark blue (loss of innervation) indicate either *acj6* mutant phenotype itself or no rescue of *acj6* mutant phenotype (*p* ≥ 0.05 comparing to *acj6* mutant innervation extent, by Chi-squared test here and after). White indicates either the wild-type innervation pattern or complete rescue of *acj6* mutant phenotype (*p* ≥ 0.05 comparing to wild-type innervation extent). Light red (partial ectopic innervation) and light blue (partial loss of innervation) indicate that the phenotypic rescue was significant (*p* < 0.05 comparing to *acj6* mutant innervation extent) but still different from wild-type (*p* < 0.05 comparing to wild-type innervation extent). Letters within the grids indicate panels where detailed quantifications are shown. Dots highlight an example of between-glomeruli additive interaction. (C–G) Examples of additive (C and D), subtractive (E), and synergistic (F and G) interactions between Acj6-regulated cell-surface proteins. (H) Summary of the cell-surface wiring executors of Acj6 for instructing correct targeting or preventing ectopic targeting of adPN dendrites to distinct glomeruli. (I) Wiring specificity of different neuron types is dictated by a cell-surface protein combinatorial code, which is controlled by combinatorial expression of transcription factors—each transcription factor regulates the expression of multiple cell-surface proteins (divergence), and each cell-surface protein is regulated by multiple transcription factors (convergence). See also Figures S7.

Notably, phenotypic rescues by different cell-surface proteins appeared to be largely non-overlapping (Figure 7A), indicating that Acj6 uses distinct cell-surface executors in different PN types for dendrite targeting.

We note that occasionally overexpression of an Acj6-regulated cell-surface protein rescued *acj6* mutant phenotypes better than overexpression of Acj6 itself (Figure 7A). Because *acj6* undergoes alternative splicing to produce 13 isoforms (Bai and Carlson, 2010; Certel *et al*., 2000), one possible explanation is that different PN types express different *acj6* isoforms to regulate the expression of different cell-surface executors, so overexpressing a specific one (isoform J in our case) may not be able to rescue all *acj6* mutant phenotypes. Another possible explanation is the delayed onset of MARCM-based overexpression due to GAL80 perdurance and the earlier action of a transcription factor compared to its downstream cell-surface executors.

Interestingly, overexpressing some cell-surface proteins that were downregulated in *acj6* mutant occasionally exacerbated dendrite mistargeting caused by *acj6* mutant. For example, the VA1v glomerulus became even less frequently innervated in *acj6^−/−^* adPN neuroblast clones overexpressing *Dg* compared to *acj6^−/−^* clones alone (Figure S7A). Examining the endogenous Dg expression using the conditional tag strategy revealed that Dg was not expressed in VA1v PNs (Figure S7B). Therefore, the worsened phenotype is likely caused by mis-expression of Dg. These results further support that Acj6 regulates different cell-surface proteins in different PN types for their precise wiring.

### Acj6 specifies PN dendrite targeting through a combinatorial cell-surface molecule code

The limited rescue of a small subset of *acj6* mutant phenotypes by each single cell-surface protein (Figure 7A) prompted us to examine whether combinatorial expression could better rescue wiring defects caused by loss of *acj6*. We tested seven combinations of two Acj6-regulated cell-surface proteins and found that these combinations indeed led to overall stronger and broader rescue of *acj6^−/−^* wiring defects in adPN neuroblast clones (more white and lighter colored squares in Figure 7B than in Figure 7A). A closer examination of these combinations revealed different modes of genetic interactions between Acj6-regulated cell-surface executors, which collectively contributed to the improved rescuing efficacy.

The most frequently observed interaction mode is additive. For each glomerulus, we observed two candidates with partial rescues could “sum up” to produce an almost complete rescue when co-expressed. For instance, overexpression of either Otk or Piezo partially restored innervation to the VM2 glomerulus (Figure 7C and ‘C’ squares in Figure 7A). Co-expression of Otk and Piezo led to wild-type-like innervation in the VM2 glomerulus (Figure 7C and the ‘C’ square in Figure 7B). When Piezo was coupled with Dg, they reduced ectopic targeting to the DL2 glomerulus from ∼80% in *acj6* mutant to less than 20% (Figure 7D and ‘D’ squares in Figures 7A and B). Across different glomeruli, the phenotypic rescue by different cell-surface executors were also additive. For example, Dop2R but not Dg overexpression rescued loss of innervation to the DA3 glomerulus, while Dg but not Dop2R overexpression rescued ectopic targeting to the VA6 glomerulus (Figure 7A, black dots). When Dop2R and Dg were co-expressed, both DA3 and VA6 phenotypes were rescued (Figure 7B, black dots). Lastly, additive interactions can inhibit phenotypic rescue if one candidate counteracts, resulting in a subtractive interaction. As described above, Dg overexpression exacerbated the loss of VA1v innervation phenotype in *acj6* mutant (Figure 7E). Co-expressing Dg with Nep3, whose expression rescued loss of innervation to the VA1v glomerulus, diminished the rescue effect of Nep3 (Figure 7E and ‘E’ squares in Figures 7A and 7B).

Besides additive interactions, we also observed synergistic interactions. For example, neither Otk nor Dop1R1 overexpression alone significantly rescued VL2a ectopic targeting in *acj6* mutant (Figure 7F and ‘F’ squares in Figure 7A). However, co-expressing both reduced the mistargeting rate from more than 70% in mutant to ∼30% (Figure 7F and ‘F’ square in Figure 7B), suggesting that they might both be required for controlling dendrite targeting. In another case, Dop2R overexpression alone could not provide significant rescue of DA4m ectopic targeting but, when co-expressed with Otk, it significantly enhanced rescue by Otk (Figure 7G and ‘G’ squares in Figures 7A and B).

Taken together, combinatorial expressions of Acj6-regulated cell-surface executors deliver stronger and broader rescues of *acj6^−/−^* wiring defects than single expressions. These results reveal that Acj6-regulated executors act combinatorically—both between different PN types and within the same PN types—to instruct dendrite targeting, and different PN types employ distinct cell-surface combinatorial codes for their precise targeting (Figure 7H).

## DISCUSSION

Nucleus-residing transcription factors control neuronal connectivity but do not directly regulate wiring at distal neurites. It is therefore a long-standing presumption that transcription factors regulate the expression of cell-surface proteins to execute wiring decisions. Here, we combined *in situ* cell-surface proteomic profiling and genetic analyses to systematically identify cell-surface executors for the lineage-specific transcription factor Acj6 in the wiring of fly olfactory PNs. We discovered many previously unknown wiring molecules, including the mechanosensitive ion channel Piezo, and revealed the operational strategies of Acj6-regulated cell-surface executors in instructing dendrite targeting.

### Identifying cell-surface executors of transcription factors

For most transcription factors directing neuronal connectivity, their cell-surface wiring executors remain elusive due to the lack of approaches for systematically identifying them. In cases where the wiring executor(s) of a transcription factor have been identified, often only one or two molecules whose wiring function has been established in that neuron type were investigated (selected examples summarized in Santiago and Bashaw, 2014), precluding the discovery of the repertoire of cell-surface wiring executors controlled by a transcription factor. RNA sequencing provides a way to determine how deleting a transcription factor alters the transcriptome of developing neurons (Jain et al., 2020; Morey et al., 2008; Peng *et al*., 2018; Peng *et al*., 2017), but studies comparing transcriptome and proteome of the same cell type often revealed modest to poor correlations (Carlyle et al., 2017; Ghazalpour et al., 2011; Wang et al., 2019), particularly for membrane proteins whose trafficking and turnover is subject to extensive post-translational regulation (Li *et al*., 2020; MacGurn et al., 2012; Trowbridge et al., 1993). Indeed, comparing wild-type and *acj6* mutant adPN transcriptomes revealed that Acj6 could regulate the expression of proteins on PN surface post-transcriptionally (Figures 1E and 1F; Figure S1).

We therefore took advantage of our recently developed cell-surface proteomic profiling (Li *et al*., 2020) to identify Acj6-regulated wiring executors for PN dendrite targeting by comparing cell-surface proteomes of wild-type and *acj6*-mutant developing PNs (Figure 2), followed by a proteome-informed genetic screen (Figure 3). Quantitative cell-surface proteomic profiling allowed us to directly examine the protein expression levels at the cell surface, instead of inferring from the mRNA levels (Aebersold and Mann, 2016; Han et al., 2018; Hosp and Mann, 2017; Li *et al*., 2020; Natividad et al., 2018). Our subsequent functional interrogations identified many Acj6 executors via both loss-of-function and rescue experiments (Figures 4–7), highlighting the effectiveness of our approach for identifying cell-surface executors of a transcription factor.

Since transcription factors often serve as central commanders of cellular functions while cell-surface proteins work as direct executors in cell-cell interactions, the ‘transcription factor → cell-surface executor → physiological function’ framework is of importance for not only neural circuit wiring but also all other biological processes involving cells communicating with their environment. Notably, cell-surface proteins are more accessible than nucleus-residing transcription factors and often exhibit higher specificity for a specific biological process than broadly-acting transcription factors, making cell-surface proteins more favorable targets for drug development. Therefore, systematically identifying cell-surface executors of a transcription factor not only delineates biological mechanisms but also has the potential for discovering more druggable targets.

### Piezo instructs neural circuit wiring independently of its mechanosensitive ion channel activity

Since the discovery of Piezo1 and Piezo2 (Coste et al., 2010), numerous functions of the Piezo protein family have been uncovered in a wide range of physiological contexts, including but not limited to somatosensory and interoceptive mechanotransduction and cell volume regulation (reviewed in Murthy et al., 2017; Szczot et al., 2021). Piezo has also been shown to regulate neurodevelopmental processes and axon regeneration—it promotes neurogenesis instead of astrogenesis in human neural stem cells (Pathak et al., 2014), contributes to axon growth and targeting of *Xenopus* retinal ganglion cells (Koser et al., 2016), and cell autonomously inhibits axon regeneration (Song et al., 2019). Notably, all known functions of Piezo thus far have been attributed to its mechanosensitive ion channel activity.

We found that Piezo instructs PN dendrite targeting, and its mechanically activated ion channel activity is dispensable for this function (Figure 5), suggesting that Piezo can also function independently of being a mechanosensitive ion channel. We note that lack of mechanically activated currents from Piezo-Myc2306 does not distinguish between loss of ion permeation property and loss of mechanosensitivity. Given that Piezo proteins form 900-kDa complexes with a large extracellular surface area, it is possible that currently unknown Piezo ligands and/or receptors mediate the wiring function of Piezo. Systematic identification of Piezo’s molecular partners may reveal how Piezo instructs dendrite targeting and how it functions independently of its mechanosensitive channel activity.

Besides Piezo, our proteome-informed genetic screen discovered several other unconventional wiring molecules for PN dendrite targeting, including the dopamine receptors Dop1R1 and Dop2R, highlighting the functional versatility of these molecules in multiple biological contexts. It will be interesting to explore whether these molecules regulate neuronal wiring via their conventional molecular functions or through currently unknown mechanisms like Piezo.

### Acj6 instructs PN wiring through a cell-surface combinatorial code

Because the number of neuronal connections far exceeds the number of wiring molecules, it has been proposed that precise neuronal connections are specified by a combinatorial code—each type of neuron uses a unique combination of wiring molecules, and each molecule is used in multiple neuron types (Hong and Luo, 2014). Here, we observed that many *acj6^−/−^* adPN dendrites ectopically targeted to glomeruli normally occupied by other (not labeled) Acj6+ adPNs (Figures 6A–6F), suggesting that Acj6 regulates the expression of distinct cell-surface proteins in different PN types and instructs dendrite targeting through a combinatorial code.

Endogenous protein tagging of three Acj6 executors, Otk, Piezo, and Dg, showed that Acj6 indeed regulates the expression of different cell-surface proteins in different PN types (Figures 4B–4D; Figures S6A and S6B; Figure S7B). Functionally, our systematic examination of Acj6 executors in PN dendrite targeting by RNAi knock-down (Figure 3; Figure S3; Figure S4) and rescue experiments (Figure 6; Figure 7; Figure S7A) revealed that different PN types require different cell-surface proteins for dendrite targeting (columns in Figure 7A), and each molecule often regulates a few distinct PN types (rows in Figure 7A). Moreover, combinatorial expression of Acj6 executors yielded stronger and broader rescues of specific *acj6* mutant phenotypes than expressing individual executors (Figure 7B). All these observations support the notion that Acj6 employs unique combinations of cell-surface proteins in different PN types to control dendrite targeting (Figure 7H).

Our findings illustrate a divergent ‘transcription factor → cell-surface executor’ relationship—one transcription factor Acj6 regulates many cell-surface proteins that execute its function (Figure 7I). This could not have been discovered in previous studies when only one or two wiring executors of a transcription factor were examined (reviewed in Santiago and Bashaw, 2014). On the other hand, Acj6-regulated cell-surface proteins often also have Acj6-independent functions. For instance, Otk is also expressed by Acj6-negative lPNs (Figures 4B–4D) and is required for lPN dendrite targeting (Figures S5H and S5I). Piezo also participates in lPN dendrite targeting (Figures S6F–S6I). Thus, they must also be regulated by other transcription factors. The fact that Acj6 differentially regulates the same cell-surface proteins in different adPN types also demands the involvement of other transcription factors; otherwise, Acj6 would uniformly regulate each executor in all Acj6+ PN types. Therefore, the ‘transcription factor → cell-surface executor’ relationship is also convergent—multiple transcription factors regulate the expression of one cell-surface protein (Figure 7I). We thus anticipate the existence of two intertwined combinatorial codes—one of the transcription factors, which specifies the other one of cell-surface executors— to determine neuronal wiring specificity.

## STAR★METHODS

- KEY RESOURCES TABLE
- RESOURCE AVAILABILITY

- Lead contact
- Material availability
- Data and code availability
- EXPERIMENTAL MODEL AND SUBJECT DETAILS

- *Drosophila* stocks and genotypes
- METHOD DETAILS

- Transcriptome profiling of adPNs
- Acj6 binding site prediction
- Biotinylation of PN-surface proteins
- Collection of biotinylated proteins
- Western blot of biotinylated proteins
- On-bead trypsin digestion of biotinylated proteins
- TMT labeling and SCX StageTip fractionation of peptides
- Liquid chromatography and mass spectrometry
- Mass spectrometry data processing
- Proteomic data cutoff analysis
- Immunocytochemistry
- Image acquisition and processing
- Genetic screen to identify molecules required for adPN dendrite targeting
- Generation of endogenous conditional tags
- Generation of UAS constructs and transgenic flies
- MARCM-based mosaic analysis
- Transfection and staining of *Drosophila* S2 cells
- Cell culture and transfection of HEK293T *PIEZO1* knockout cells
- Electrophysiology
- QUANTIFICATION AND STATISTICAL ANALYSIS

- RNA-seq data analysis
- Quantitative comparison of wild-type and *acj6* mutant proteomes
- Quantification of Otk and Piezo expression in PNs
- Quantification of adPN dendrite innervation patterns in MARCM-based mosaic analysis

## ACKNOWLEDGMENTS

We thank Y. Rao, V. Ruta, Y. Aso, the Bloomington *Drosophila* Stock Center, and the Vienna *Drosophila* Resource Center for the fly lines, and A. Patapoutian and Addgene for plasmids. We thank T. Clandinin, J. Kaltschmidt, K. Shen, C. McLaughlin, Z. Li, K. Wong, C. Lyu, Y. Ge, J. Ren, D. Pederick, M. Wagner, and members of the Luo lab for technical support and/or insightful advice on this study. We thank M. Molacavage for administrative assistance. Q.X. was a Bertarelli Fellow. J.L. is a Jane Coffin Childs postdoctoral fellow. L.L. is an investigator of the Howard Hughes Medical Institute. This work was supported by the National Institutes of Health (R01-DC005982 to L.L. and R01-DK121409 to S.A.C. and A.Y.T.) and the Wu Tsai Neurosciences Institute of Stanford University (Neuro-omics grant to A.Y.T., S.R.Q., and L.L.).

## AUTHOR CONTRIBUTIONS

Q.X., J.L., and L.L. conceived this project. Q.X., J.L., and L.L. designed the proteomic experiments with input from S.H. and A.Y.T. Q.X., J.L., and S.A.S. collected pupae and Q.X., J.L., H.L., C.X., T.L., and D.J.L. dissected fly brains for the proteomic experiment. J.L. and S.H. processed proteomic samples. C.X. hand-delivered proteomic samples to the Broad Institute after FedEx messed up the shipment. N.D.U., T.S., D.R.M., and S.A.C. performed post-enrichment sample processing, mass spectrometry, and initial data analysis. Q.X., J.L., and S.H. analyzed proteomic data. D.O. performed and analyzed the electrophysiology data under S.E.M.’s guidance. Q.X. and H.L. performed transcriptome experiments and analysis with the support of S.R.Q. Q.X. designed and performed all other experiments with input from J.L. and L.L. and assistance from S.K., R.G., W.W., and D.J.L. Q.X., J.L., and L.L. wrote the manuscript with input from all authors. L.L supervised the project.

## STAR★METHODS

### KEY RESOURCES TABLE

| REAGENT or RESOURCE | SOURCE | IDENTIFIER |
| --- | --- | --- |
| Antibodies |  |  |
| Rat anti-NCad | Developmental Studies Hybridoma Bank | RRID: AB_528121 |
| Chicken anti-GFP | Aves Labs | RRID: AB_10000240 |
| Mouse anti-FLAG | Sigma-Aldrich | RRID: AB_259529 |
| Mouse anti-V5 | Sigma-Aldrich | RRID: AB_2556564 |
| Deposited data |  |  |
| RNA sequencing data | This paper | Will be deposited to GEO |
| Mass spectrometry proteomics data | This paper | Will be deposited to MassIVE |
| Experimental models: Organisms/strains |  |  |
| *D. melanogaster: tubP-GAL80* | (Lee and Luo, 1999) | RRID: BDSC_9917 |
| *D. melanogaster: UAS-mCD8-GFP* | (Lee and Luo, 1999) | RRID: DGRC_108068 |
| *D. melanogaster: hsFlp* | (Golic and Lindquist, 1989) | N/A |
| *D. melanogaster: FRT19A* | (Xu and Rubin, 1993) | RRID: BDSC_1744 |
| *D. melanogaster: FRT40A* | (Xu and Rubin, 1993) | N/A |
| *D. melanogaster: FRT42D* | (Xu and Rubin, 1993) | RRID: BDSC_1802 |
| *D. melanogaster: GH146-Flp* | (Hong *et al.*, 2009) | N/A |
| *D. melanogaster: acj6<sup>6</sup>* | (Clyne *et al.*, 1999) | RRID: BDSC_60686 |
| *D. melanogaster: otk<sup>3</sup>* | (Winberg *et al.*, 2001) | N/A |
| *D. melanogaster: piezo<sup>KO</sup>* | (Kim *et al.*, 2012) | RRID: BDSC_58770 |
| *D. melanogaster: VT033006-GAL4* | (Tirian and Dickson, 2017) | RRID: VDRC_202281 |
| *D. melanogaster: GH146-GAL4* | (Stocker *et al.*, 1997) | RRID: BDSC_30026 |
| *D. melanogaster: VT033006-GAL4<sup>DBD</sup>* | Yoshi Aso (unpublished) | N/A |
| *D. melanogaster: C15-p65<sup>AD</sup>* | (Xie *et al.*, 2019) | N/A |
| *D. melanogaster: UAS-HRP-CD2* | (Larsen *et al.*, 2003) | RRID: BDSC_9906 |
| *D. melanogaster: UAS-RNAi lines* | (Dietzl *et al.*, 2007; Ni *et al.*, 2011; Perkins *et al.*, 2015) | Stock numbers listed in Figure S3 |
| *D. melanogaster: UAS-dcr2* | (Dietzl *et al.*, 2007) | N/A |
| *D. melanogaster: UAS-otk* | (Winberg *et al.*, 2001) | N/A |
| *D. melanogaster: UAS-piezo* | This study | N/A |
| *D. melanogaster: UAS-piezo-2306Myc* | This study | N/A |
| *D. melanogaster: UAS-acj6-J* | (Bai and Carlson, 2010) | N/A |
| *D. melanogaster: UAS-Dg* | Robert Ray, Francis Crick Institute | RRID: BDSC_63052 |
| *D. melanogaster: UAS-Dop1R1* | This study | N/A |
| *D. melanogaster: UAS-Dop2R* | (Deng *et al.*, 2019) | RRID: BDSC_78798 |
| *D. melanogaster: UAS-Nep3* | This study | N/A |
| *D. melanogaster: UAS-CG5027* | This study | N/A |
| *D. melanogaster: UAS-Osty* | This study | N/A |
| *D. melanogaster: otk-FRT-V5-STOP-FRT-FLAG-STOP* | This study | N/A |
| *D. melanogaster: piezo-FRT-V5-STOP-FRT-FLAG-STOP* | This study | N/A |
| *D. melanogaster: Dg-FRT-V5-STOP-FRT-FLAG-STOP* | This study | N/A |
| HEK-PI KO cells | (Lukacs *et al.*, 2015) | N/A |

| Recombinant DNA |  |  |
| --- | --- | --- |
| Conditional tag cassette *FRT-1xV5-6xStop-loxP-mini-White-loxP-FRT-3xFLAG-6xStop* | (Li *et al.*, 2020) | N/A |
| *pCR-Blunt-TOPO* | Thermo Fisher Scientific | Catalog #: K280020 |
| *pU6-BbsI-chiRNA* | (Gratz *et al.*, 2014; Gratz *et al.*, 2013) | RRID: Addgene_45946 |
| *pUASTattB* | (Bischof *et al.*, 2007) | RRID: DGRC_1419 |
| *pUAST-piezo-901Myc-FLAG* | This study | N/A |
| *pUAST-piezo-2289Myc-FLAG* | This study | N/A |
| *pUAST-piezo-2291Myc-FLAG* | This study | N/A |
| *pUAST-piezo-2306Myc-FLAG* | This study | N/A |
| *pcDNA3.2-piezo-IRES2-mNeonGreen* | Ardem Patapoutian, Scripps Research Institute | N/A |
| *pcDNA3.2-piezo-WT-IRES2-mNeonGreen* | This study | N/A |
| *pcDNA3.2-piezo-2289Myc-mNeonGreen* | This study | N/A |
| *pcDNA3.2-piezo-2291Myc-mNeonGreen* | This study | N/A |
| *pcDNA3.2-piezo-2306Myc-mNeonGreen* | This study | N/A |
| Software and algorithms |  |  |
| Zen | Carl Zeiss | RRID: SCR_013672 |
| ImageJ | National Institutes of Health | RRID: SCR_003070 |
| Illustrator | Adobe | RRID: SCR_010279 |
| Spectrum Mill | Agilent | https://www.agilent.com/Agilent404?s=https://www.agilent.com/en/products/software-informatics/masshunter-suite/masshunter-for-life-science-research/spectrum-mill |
| pClamp | Molecular Devices | RRID: SCR_011323 |

### RESOURCE AVAILABILITY

#### Lead contact

Further information and requests for resources and reagents should be directed to the Lead Contact, Liqun Luo.

#### Material availability

All unique reagents generated in this study are available from the Lead Contact.

#### Data and code availability

Raw sequencing reads will be deposited to GEO. Processed RNA sequencing data is provided in Table S1. Mass spectrometry proteomics data will be deposited to MassIVE. Original and processed proteomic data is provided in Table S2. Custom analysis code will be made available at https://github.com/Qijing-Xie/Acj6.

### EXPERIMENTAL MODEL AND SUBJECT DETAILS

#### *Drosophila* stocks and genotypes

Flies were maintained on standard cornmeal medium with a 12 hr light–dark cycle at 25°C, except for the RNAi/overexpressing screen in which flies were raised at 29°C. Complete genotypes of flies used in each experiment are described in Table S3. The following lines were used in this study: *tubP-GAL80* and *UAS-mCD8-GFP* (Lee and Luo, 1999), *hsFlp* (Golic and Lindquist, 1989), *FRT19A*, *FRT40A*, and *FRT42D* (Xu and Rubin, 1993), *GH146-Flp* (Hong *et al*., 2009), *acj6^6^* (Clyne *et al*., 1999), *otk^3^* (Winberg *et al*., 2001), *piezo^KO^* (Kim *et al*., 2012), *VT033006-GAL4* (Tirian and Dickson, 2017), *GH146-GAL4* (Stocker *et al*., 1997), *C15-p65^AD^* (Xie *et al*., 2019), *UAS-HRP-CD2* (Larsen *et al*., 2003), *UAS-dcr2* (Dietzl *et al*., 2007), *UAS-otk* (Winberg *et al*., 2001), *UAS-acj6-J* (Bai and Carlson, 2010). *UAS-Dg* (Robert Ray, Francis Crick Institute), and *UAS-Dop2R* (Deng *et al*., 2019). The RNAi lines were generated previously (Dietzl *et al*., 2007; Ni *et al*., 2011; Perkins *et al*., 2015) and acquired from the Bloomington Drosophila Stock Center and the Vienna Drosophila Resource Center. *VT033006-GAL4^DBD^* is unpublished reagent generously provided by Yoshi Aso (Janelia Research Campus). *UAS-piezo*, *UAS-piezo-2306Myc*, *UAS-Dop1R1*, *UAS-Nep3*, *UAS-CG5027*, *UAS-Ostγ*, *otk-FRT-V5-STOP-FRT-FLAG-STOP*, *piezo-FRT-V5-STOP-FRT-FLAG-STOP*, and *Dg-FRT-V5-STOP-FRT-FLAG-STOP* were generated in this study.

### METHOD DETAILS

#### Transcriptome profiling of adPNs

Wild-type or *acj6* mutant *Drosophila* brains with mCD8-GFP labeled adPNs using the split GAL4 driver *C15-p65^AD^, VT033006-GAL4^DBD^* were dissected at 36–40 hr after puparium formation in Schneider’s medium (Thermo Fisher). Single-cell suspension was prepared as previously described (Li et al., 2017). One thousand GFP positive cells were sorted for each sample using Fluorescence Activated Cell Sorting (FACS) on a Sony SH800 cell sorter system (Sony Biotechnology). Three biological replicates were collected for each genotype. Full-length poly(A)- tailed RNA was reverse-transcribed and amplified by PCR following the modified SMART-seq2 protocol (Picelli et al., 2014). Sequencing libraries were prepared from amplified cDNA, pooled, and quantified using BioAnalyser (Agilent). Sequencing was performed using the NextSeq 500 Sequencing system (Illumina) with 75 paired-end reads.

#### Acj6 binding site prediction

Prediction of Acj6 binding site was adapted from previously described method with experimentally validated Acj6 consensus (Bai et al., 2009). For each gene, we scanned through its promoter sequence, encompassing sequence from 500bp upstream to 100bp downstream of its transcription start site (Eukaryotic Promoter Database; Dreos et al., 2017; Dreos et al., 2015). The sum of weight calculated using the positional weight matrix (PWM) was used to score each of the 12bp long DNA sequence on both the sense and antisense strand. Sequences with a score greater than 6 were counted as putative Acj6 binding sites.

#### Biotinylation of PN-surface proteins

PN surface biotinylation was performed following the previously published method (Li *et al*., 2020). Briefly, we dissected wild-type or *acj6* mutant brains expressing HRP-rCD2 by PNs in pre-chilled Schneider’s medium (Thermo Fisher), removed the optic lobes, and transferred them into 500 μL of the Schneider’s medium in 1.5 mL protein low-binding tubes (Eppendorf) on ice. Brains were washed with fresh Schneider’s medium to remove fat bodies and debris and were incubated in 100 μM of BxxP in Schneider’s medium on ice for one hour. Brains were then labeled with 1 mM (0.003%) H_2_O_2_ (Thermo Fisher) for 5 minutes, and immediately quenched by five thorough washes using quenching buffer (10 mM sodium ascorbate [Spectrum Chemicals], 5 mM Trolox [Sigma-Aldrich], and 10 mM sodium azide [Sigma-Aldrich] in phosphate buffered saline [PBS; Thermo Fisher]). After the washes, the quenching solution was removed, and brains were either fixed for immunostaining (see below for details) or were snapped frozen in liquid nitrogen and stored at –80°C for the proteomic sample collection. For the proteomic sample collection, 1100 dissected and biotinylated brains were collected for each experimental group (8800 brains in total).

#### Collection of biotinylated proteins

For each proteomic sample, there were five tubes each containing ∼220 dissected brains. We added 40 μL of high-SDS RIPA (50mM Tris-HCl [pH 8.0], 150 mM NaCl, 1% sodium dodecyl sulfate [SDS], 0.5% sodium deoxycholate, 1% Triton X-100, 1x protease inhibitor cocktail [Sigma-Aldrich; catalog # P8849], and 1 mM phenylmethylsulfonyl fluoride [PMSF; Sigma-Aldrich]) to each of those tubes, and grinded the samples on ice using disposable pestles with an electric pellet pestle driver. Tubes containing brain lysates of the same experimental group were spun down, merged, and rinsed with an additional 100 μL of high-SDS RIPA to collect remaining proteins. Samples were then vortexed briefly, sonicated twice for ten seconds each, and incubated at 95°C for five minutes to denature postsynaptic density proteins. 1.2 mL of SDS-free RIPA buffer (50 mM Tris-HCl [pH 8.0], 150 mM NaCl, 0.5% sodium deoxycholate, 1% Triton X-100, 1x protease inhibitor cocktail and 1 mM PMSF) were added to each sample, and the mixture was rotated for two hours at 4°C. Lysates were then diluted with 200 μL of normal RIPA buffer (50 mM Tris-HCl [pH 8.0], 150 mM NaCl, 0.2% SDS, 0.5% sodium deoxycholate, 1% Triton X-100, 1x protease inhibitor cocktail, and 1 mM PMSF), transferred to 3.5 mL ultracentrifuge tubes (Beckman Coulter), and centrifuged at 100,000 g for 30 minutes at 4°C. 1.5 mL of the supernatant was carefully collected for each sample.

400 μL of streptavidin-coated magnetic beads (Pierce; catalog # 88817) washed twice using 1 ml RIPA buffer were added to each of the post-ultracentrifugation brain lysates. The lysate and the streptavidin bead mixture were left to rotate at 4°C overnight. On the following day, beads were washed twice with 1 mL RIPA buffer, once with 1 mL of 1 M KCl, once with 1 mL of 0.1 M Na_2_CO_3_, once with 1 mL of 2 M urea in 10 mM Tris-HCl (pH 8.0), and again twice with 1 mL RIPA buffer. The beads were resuspended in 1 mL fresh RIPA buffer. 35 μL of the bead suspension was taken out for western blot, and the rest proceeded to on-bead digestion.

#### Western blot of biotinylated proteins

Biotinylated proteins were eluted from streptavidin beads by the addition of 20 μL elution buffer (2X Laemmli sample buffer, 20 mM DTT, 2 mM biotin) followed by a 10 min incubation at 95°C. Proteins were resolved by 4%–12% Bis-Tris PAGE gels (Thermo Fisher) and transferred to nitrocellulose membrane (Thermo Fisher). After blocking with 3% bovine serum albumin (BSA) in Tris-buffered saline with 0.1% Tween 20 (TBST; Thermo Fisher) for 1 hour, membrane was incubated with 0.3 mg/mL HRP-conjugated streptavidin for one hour. The Clarity Western ECL blotting substrate (Bio-Rad) and BioSpectrum imaging system (UVP) were used to develop and detect chemiluminescence.

#### On-bead trypsin digestion of biotinylated proteins

To prepare proteomic samples for mass spectrometry analysis, proteins bound to streptavidin beads were washed twice with 200 μL of 50 mM Tris-HCl buffer (pH 7.5) and twice with 2 M urea/50 mM Tris (pH 7.5) buffer. After washes, the 2 M urea/50 mM Tris (pH 7.5) buffer was removed, and beads were incubated with 80 μL of 2 M urea/50 mM Tris buffer containing 1 mM dithiothreitol (DTT) and 0.4 μg trypsin for 1 hour at 25°C with shaking at 1000 revolutions per minute (rpm). After 1 hour, the supernatant was transferred to a fresh tube. The streptavidin beads were rinsed twice with 60 μL of 2 M urea/50 mM Tris (pH 7.5) buffer and the solution was combined with the on-bead digest supernatant. The eluate was reduced with 4 mM DTT for 30 minutes at 25°C with shaking at 1000 rpm and alkylated with 10 mM iodoacetamide for 45 minutes in the dark at 25°C while shaking at 1000 rpm. An additional 0.5 μg of trypsin was added to the sample and the digestion was completed overnight at 25°C with shaking at 700 rpm. After overnight digestion, the sample was acidified (pH < 3) by adding formic acid (FA) such that the sample contained 1% FA. Samples were desalted on C18 StageTips (3M). Briefly, C18 StageTips were conditioned with 100 μL of 100% MeOH, 100 μL of 50% MeCN/0.1% FA followed by two washes with 100μL of 0.1% FA. Acidified peptides were loaded onto the conditioned StageTips, which were subsequently washed twice with 100 μL of 0.1% FA. Peptides were eluted from StageTips with 50 μL of 50% MeCN/0.1 % FA and dried to completion.

#### TMT labeling and SCX StageTip fractionation of peptides

8 TMT reagents from a 10-plex reagent kit were used to label desalted peptides (Thermo Fisher) as directed by the manufacturer. Peptides were reconstituted in 100 μL of 50 mM HEPES. Each 0.8 mg vial of TMT reagent was reconstituted in 41 μL of anhydrous acetonitrile and incubated with the corresponding peptide sample for 1 hour at room temperature. Labeling of samples with TMT reagents was completed with the design described in Figure 2A. TMT labeling reactions were quenched with 8 μL of 5% hydroxylamine at room temperature for 15 minutes with shaking, evaporated to dryness in a vacuum concentrator, and desalted on C18 StageTips as described above. For the TMT 8-plex cassette, 50% of the sample was fractionated into 3 fractions by Strong Cation Exchange (SCX) StageTips while the other 50% of each sample was reserved for LC-MS analysis by a single shot, long gradient. One SCX StageTip was prepared per sample using 3 plugs of SCX material (3M) topped with 2 plugs of C18 material. StageTips were sequentially conditioned with 100 μL of MeOH, 100 μL of 80% MeCN/0.5% acetic acid, 100 μL of 0.5% acetic acid, 100 μL of 0.5% acetic acid/500mM NH_4_AcO/20% MeCN, followed by another 100 μL of 0.5% acetic acid. Dried sample was re-suspended in 250 μL of 0.5% acetic acid, loaded onto the StageTips, and washed twice with 100 μL of 0.5% acetic acid. Sample was trans-eluted from C18 material onto the SCX with 100 μL of 80% MeCN/0.5% acetic acid, and consecutively eluted using 3 buffers with increasing pH—pH 5.15 (50mM NH_4_AcO/20% MeCN), pH 8.25 (50mM NH_4_HCO_3_/20% MeCN), and finally pH 10.3 (0.1% NH_4_OH, 20% MeCN). Three eluted fractions were re-suspended in 200 μL of 0.5% acetic acid to reduce the MeCN concentration and subsequently desalted on C18 StageTips as described above. Desalted peptides were dried to completion.

#### Liquid chromatography and mass spectrometry

Desalted, TMT-labeled peptides were resuspended in 9 μL of 3% MeCN, 0.1% FA and analyzed by online nanoflow liquid chromatography tandem mass spectrometry (LC-MS/MS) using a Q Exactive HF-X (Thermo Fisher) coupled on-line to a Proxeon Easy-nLC 1000 (Thermo Fisher). 4 μL of each sample was loaded at 500 nL/min onto a microcapillary column (360 μm outer diameter x 75 μm inner diameter) containing an integrated electrospray emitter tip (10 μm), packed to approximately 24 cm with ReproSil-Pur C18-AQ 1.9 μm beads (Dr. Maisch GmbH) and heated to 50°C. The HPLC solvent A was 3% MeCN, 0.1% FA, and the solvent B was 90% MeCN, 0.1% FA. Peptides were eluted into the mass spectrometer at a flow rate of 200 nL/min. Non-fractionated samples were analyzed using a 260 min LC-MS/MS method with the following gradient profile: (min:%B) 0:2; 1:6; 235:30; 244:60; 245:90; 250:90; 251:50; 260:50 (the last two steps at 500 nL/min flow rate). The SCX fractions were run with 110-minute method, which used the following gradient profile: (min:%B) 0:2; 1:6; 85:30; 94:60; 95:90;100:90; 101:50; 110:50 (the last two steps at 500 nL/min flow rate). The Q Exactive HF-X was operated in the data-dependent mode acquiring HCD MS/MS scans (r = 45,000) after each MS1 scan (r = 60,000) on the top 20 most abundant ions using an MS1 target of 3E6 and an MS2 target of 5E4. The maximum ion time utilized for MS/MS scans was 120 ms (single-shot) and 105 ms (SCX fractions); the HCD normalized collision energy was set to 31; the dynamic exclusion time was set to 20 s, and the peptide match and isotope exclusion functions were enabled. Charge exclusion was enabled for charge states that were unassigned, 1 and > 7.

#### Mass spectrometry data processing

Collected data were analyzed using the Spectrum Mill software package v6.1 pre-release (Agilent Technologies). Nearby MS scans with the similar precursor m/z were merged if they were within ± 60 s retention time and ± 1.4 m/z tolerance. MS/MS spectra were excluded from searching if they failed the quality filter by not having a sequence tag length 0 or did not have a precursor MH+ in the range of 750 – 4000. All extracted spectra were searched against a UniProt database containing *Drosophila melanogaster* reference proteome sequences. Search parameters included: ESI QEXACTIVE-HCD-v2 scoring parent and fragment mass tolerance of 20 ppm, 30% minimum matched peak intensity, trypsin allow P enzyme specificity with up to four missed cleavages, and calculate reversed database scores enabled. Fixed modifications were carbamidomethylation at cysteine. TMT labeling was required at lysine, but peptide N termini were allowed to be either labeled or unlabeled. Allowed variable modifications were protein N-terminal acetylation and oxidized methionine. Individual spectra were automatically assigned a confidence score using the Spectrum Mill auto-validation module. Score at the peptide mode was based on target-decoy false discovery rate (FDR) of 1%. Protein polishing auto-validation was then applied using an auto thresholding strategy. Relative abundances of proteins were determined using TMT reporter ion intensity ratios from each MS/MS spectrum and the median ratio was calculated from all MS/MS spectra contributing to a protein subgroup. Proteins identified by 2 or more distinct peptides and ratio counts were considered for the dataset.

#### Proteomic data cutoff analysis

We used a ratiometric strategy (Hung *et al*., 2014) to remove contaminants. Briefly, all detected proteins (2332 with 2 or more unique peptides) were annotated as either true-positives (TPs; proteins with plasma membrane annotation), false-positives (FPs; proteins with either cytosol, mitochondrion, or nucleus annotation but without the membrane annotation), or other annotations according to the subcellular localization annotation in the UniProt database. For each experimental group, we calculated the TMT ratios of proteins in this group compared to one of the controls and sorted proteins in a descending order. For each TMT ratio, a true-positive rate (TPR) and a false positive rate (FPR) were calculated by summing up the number of TPs or FPs with a higher ranking and divided them by the total number of TPs or FPs, respectively. The TPRs and FPRs were used to generate the ROC curves. The cutoff for each experimental-to-control group was determined by finding the TMT ratio where [TPR – FPR] is maximized. Proteins with TMT ratio higher than the cutoff were retained for each experimental group. 459 proteins that passed the cutoffs using both 127N/127C and 126/128N ratios were retained for wild-type, and 537 proteins that passed the cutoffs using both 129N/129C and 128C/130N ratios were retained for *acj6* mutant in Figure 2. We also tested a more stringent cutoff criterion where we compared each of the experimental group to both controls for each genotype, and only kept proteins that passed the cutoffs in all four possible combinations (Figures S2D–S2F). Gene Ontology analyses were performed on these gene sets using Flymine (Lyne et al., 2007).

#### Immunocytochemistry

Fly brains were dissected and immunostained according to the previously published protocol (Wu and Luo, 2006a). Briefly, fly brains of the desired genotype and developmental stage were dissected in PBS, transferred to a tube with 1 mL of fixation buffer (4% paraformaldehyde in PBS with 0.006% Triton X-100) on ice, and fixed for 20 minutes by nutating at room temperature. After fixation, brains were washed with PBST (0.3% Triton X-100 in PBS) twice, nutated in PBST for 20 minutes twice, and blocked in 5% normal goat serum in PBST for 1 hour (except for conditional tag flies; see below for details). To visualize the antennal lobe glomeruli and PN dendrites, brains were then incubated in rat anti-Ncad (N-Ex #8; 1:40; Developmental Studies Hybridoma Bank) and chicken anti-GFP (1:1000; Aves Labs) diluted in 5% normal goat serum in PBST for two overnights on a 4°C nutator. After primary antibody incubation, brains were washed four time with PBST (two quick washes and two 20-minute washes) and incubated with secondary antibodies conjugated to Alexa Fluor 488 and 647 (1:250 in 5% normal goat serum; Jackson ImmunoResearch). To visualize biotinylated proteins, brains were incubated with Neutravidin (Thermo Fisher) pre-conjugated to Alexa Fluor 647 (Invitrogen). After the antibody incubation(s), brains were washed four times (two quick washes and two 20-minute washes), mounted with SlowFade antifade reagent (Thermo Fisher), and stored at 4°C before imaging.

For the staining of Otk, Piezo, or Dg conditional tag, the above method produced low signal-to-noise FLAG (or V5) signal, so Alexa 488 Tyramide SuperBoost kit (Thermo Fisher) was used to amplify the signal. Briefly, after dissection, fixation, and washing steps, brains were rinsed with PBS twice and incubated with 3% hydrogen peroxide for 1 hour to quench the activity of endogenous peroxidases. Brains were then washed with PBST three times, blocked for 1 hour in 10% goat serum provided by the kit, and incubated with V5 or FLAG antibody diluted in 10% goat serum for two overnights on a 4°C nutator. After four 20-minute washes using PBST, brains were incubated with the poly-HRP-conjugated secondary antibody provided in the kit for two overnights on a 4°C nutator. Brains were washed four times again with PBST (two quick and two 20-minute washes) and twice with PBS. Afterwards, the brains were incubated with the tyramide solution for 5 minutes at room temperature, and reaction was immediately quenched by three washes using the quenching buffer provided by the kit. Brains were then washed with PBST four times, and NCad staining was performed using standard immunostainng protocol described above.

#### Image acquisition and processing

Images were obtained with a Zeiss LSM 780 laser-scanning confocal microscope (Carl Zeiss) using a 40x oil objective. Z-stacks were acquired at 1-μm intervals at the resolution of 512x512. Brightness and contrast adjustments as well as image cropping were done using ImageJ.

#### Genetic screen to identify molecules required for adPN dendrite targeting

The adPN screening line was generated by recombining *UAS-dcr2* with *UAS-mCD8-GFP* on the X chromosome and *C15-p65^AD^* with *VT033006-GAL4^DB^* on the third chromosome. Virgin female flies from this screening line were crossed to *UAS-RNAi* or *UAS-cDNA* males, and the progenies were kept at 25°C for 2 days after egg laying and then transferred to 29°C to enhance expression by the GAL4/UAS system. Brains were dissected, immunostained, and imaged as described above.

To quantify adPN dendrite innervation pattern, individual glomeruli were identified using the NCad staining (based on their stereotyped shapes and positions), and then categorical innervation scores (not innervated, weakly innervated, or strongly innervated) were assigned to each of the identified glomeruli. The genotypes were blinded during scoring. Data was analyzed using python packages Pandas and Scipy. For each glomerulus, we calculated the frequency of each type of innervation and plotted the results as stacked bars in Figure S3. To quantify if knocking down or overexpressing a gene caused significant dendrite innervation changes, Chi-squared tests were performed on the innervation degree frequencies of a glomerulus of a given genotype compared to control. *p-*values were adjusted for multiple comparisons using Bonferroni correction, and glomeruli whose dendrite innervation were significantly changed (*p-*value < 0.05) were color blue or red in a heatmap shown in Figure S4.

#### Generation of endogenous conditional tags

Otk, Piezo, or Dg conditional tag flies were generated based on the previously described method (Li *et al*., 2020). To make the homology-directed repair (HDR) vectors, a ∼2000bp of genomic sequence flanking the stop codon was amplified using Q5 hot-start high-fidelity DNA polymerase (New England Biolabs) and inserted into *pCR-Blunt-TOPO* vector (Thermo Fisher). The conditional tag cassette (*FRT-1xV5-6xStop-loxP-mini-White-loxP-FRT-3xFLAG-6xStop*) was amplified from the LRP1 plasmid (Li *et al*., 2020) and inserted into the *TOPO genomic sequence plasmid* to replace the stop coding using NEBuilder HiFi DNA assembly master mix (New England Biolabs). CRIPSR guide RNA (gRNA) targeting a locus near the stop codon was designed using the flyCRISPR Target Finder tool and cloned into the *pU6-BbsI-chiRNA* vector (Addgene #45946) by NEBuilder HiFi DNA assembly master mix.

The HDR and the gRNA vectors were co-injected into *vas-Cas9* (Port et al., 2014) fly embryos. G_0_ flies were crossed to a *white–* balancer and all *white+* progenies were individually balanced. To remove the *loxP-*flanked *miniWhite* cassette, each line was crossed to flies with *hs-Cre*. Fertilized eggs or young larvae were heat shocked twice at 37°C for 1 hour separated a day apart and crossed to a balancer. Their *white–* progenies were individually balanced and verified by sequencing to obtain *gene-FRT-V5-STOP-FRT-FLAG-STOP*.

#### Generation of UAS constructs and transgenic flies

To generate UAS flies, we used Q5 hot-start high-fidelity DNA polymerase (New England Biolabs) to amplify the transcripts of each candidate gene from complementary DNA (cDNA) synthesized using the total RNA of *w^1118^* fly heads (extracted using MiniPrep kit; Zymo Research, R1054). For each candidate gene, we designed more than 2 pairs of PCR primers in the 5’ and 3’ untranslated regions and inserted all resulting PCR products into *pCR-Blunt-TOPO* vector (Thermo Fisher). The *TOPO-transcript* vectors of the same gene were sequenced and compared to verify that no error was introduced to the coding sequence during reverse transcription. The verified coding sequences were then amplified and assembled into either the *pUAST-attB* vector (*Nep3*) or a modified *pUAST-attB* vector in which a FLAG tag was added at the 3’ end (*Dg*, *CG5027*, *DopR*, *Ostγ*, and *piezo*). To generate Piezo mutants, myc tag was inserted into amino acid position 901, 2289, 2291, and 2306 (Figure 5; Figure S6) in plasmid *pUAST-attB-piezo-FLAG* using Q5 site-directed mutagenesis kit (New England Biolabs). Each plasmid was sequence confirmed by full-length DNA sequencing.

The *pUAST-attB* constructs were inserted into either *attP40* or *attP86Fb* (for *Piezo* constructs) landing sites. G_0_ flies were crossed to a *white^−^* balancer, and all *white^+^* progenies were individually balanced and verified by sequencing.

#### MARCM-based mosaic analysis

*hsFlp*-based MARCM analyses were performed following the previously published protocol (Wu and Luo, 2006b). Each fly contains *GH146-GAL4*, *UAS-mCD8-GFP*, *tubP-GAL80*, *hsFlp*, the desired *FRT*, a mutant allele distal to the *FRT* site (or wild-type for control), and, in rescue experiments, one or two *UAS-candidate gene* (see Table S3 for complete genotypes). To generate adPN neuroblast, lPN neuroblast, or DL1 single-cell MARCM clones, flies were heat shocked at 0–24 hours after larvae hatching for 1 hour at 37°C. More than 150 adult brains were dissected for each genotype to get approximately 20 clones.

#### Transfection and staining of *Drosophila* S2 cells

*pUAST-attB-piezo-Myc-FLAG* constructs contain an intracellular C-terminal FLAG tag and an extracellular Myc tag inserted at different positions. Piezo-901Myc, the equivalent of mPiezo1- 897Myc, was used as a control to show wild-type Piezo expression (Coste *et al*., 2015; Saotome et al., 2018). S2 cells were co-transfected with these Piezo constructs and *Actin-GAL4* using FuGENE HD transfection Reagent (Promega). After 48 hours, transfected cells were incubated with rabbit anti-Myc antibody (1:250; Cell Signaling) and mouse anti-FLAG M2 antibody (1:200; Sigma-Aldrich) either before or after 4% PFA fixation and permeabilization with 0.3% Triton in PBS for non-permeabilized and permeabilized condition, respectively. Cells were then washed with PBS, incubated with secondary antibodies, and imaged.

#### Cell culture and transfection of HEK293T *PIEZO1* knockout cells

Human embryonic kidney HEK293T *PIEZO1* knockout (HEK-P1KO) cells were grown in Dulbecco’s Modified Eagle Medium (DMEM) with 10% fetal bovine serum, and 1% penicillin and streptomycin. Cells were plated on coverslips coated with laminin and transfected with 700 ng of plasmid DNA using lipofectamine 2000 (Thermo Fisher Scientific, Cat No. 11668019) as per manufacturer’s instructions. The cDNA sequence of wild-type and mutant *Drosophila* Piezos were cloned into pcDNA3.2 vector that has been modified to include an IRES2-mNeonGreen element.

#### Electrophysiology

Cells were recorded from 48–72 hours following transfection at room temperature. All recordings were done in whole-cell voltage clamp mode using Axopatch 200B amplifier (Axon Instruments). Currents were acquired at membrane holding potential of –80mV, sampled at 20 kHz and filtered at 2 kHz. Leak currents before mechanical stimulations were subtracted off-line from the current traces in Clampex 11. Current traces were analyzed using Clampex 11 and GraphPad Prism.

Extracellular solution was composed of: 133 mM NaCl, 3 mM KCl, 2.5 mM CaCl_2_, 1 mM MgCl_2_, 10 mM HEPES, and 10 mM Glucose, 310 mOsm/L, and pH of 7.3 (NaOH). Intracellular solution was composed of: 133mM CsCl, 5mM EGTA, 1 mM CaCl_2_, 1 mM MgCl_2_, 10 mM HEPES, 4 mM Mg_2_ATP, and 0.4 mM Na_2_GTP, 300 mOsm/L, and pH of 7.3 (CsOH). Patch pipettes were made from borosilicate capillaries pulled with Sutter Instruments puller (Model P- 97). Pipettes of 1.8–4 MΩ resistance in intracellular solution were used for recordings.

Mechanical stimulation was induced using a blunt glass probe held at ∼80° angle, and controlled by a piezoelectric actuator (E625 LVPZT, Physik Instrumente). Cells were indented for 300 ms at 1-micron increments, with a 30 s inter-sweep duration. Seal integrity was continuously monitored by including a 10 ms voltage step from –80 mV to –75 mV, 90 ms prior to indenting the cell.

### QUANTIFICATION AND STATISTICAL ANALYSIS

The statistical tests and numbers of independent replicates per experiment are indicated in the figures or figure legends.

#### RNA-seq data analyses

Reads were aligned to the *Drosophila melanogaster* genome (r6.10) using STAR (2.4.2) (Dobin et al., 2013). Gene counts were produced using HTseq (0.7.2) with default settings except ‘‘-m intersection-strict’ (Anders et al., 2015). To normalize for differences in sequencing depth across individual cells, we rescaled gene counts to transcript counts per million reads (CPM). Genes with less than 1 CPM in more than 3 samples were excluded from further analysis.

Differential gene expression was calculated with R package edgeR (Robinson et al., 2010) and limma (Ritchie et al., 2015). Briefly, data were normalized with the weighted trimmed mean of M-values (TMM) method (Robinson and Oshlack, 2010) and fitted to a linear model with voom and lmFit functions (Ritchie *et al*., 2015). Finally, empirical Bayes statistics were applied to correct variance of genes with low expression. *p*-value was adjusted using the Benjamini–Hochberg method. We define differentially expressed genes to be those with adjusted *p*-value less than 0.05. Enrichment analysis was performed using Enrichr (Kuleshov et al., 2016) and pathway gene sets “KEGG_2019” in the GSEApy python package.

#### Quantitative comparison of wild-type and *acj6* mutant proteomes

For the volcano plots (Figures 2G; Figure S2F) comparing differentially enriched proteins on *acj6* mutant PN surface compared to wild-type PN surface, a linear model was fit to account for the variance across replicates for each stage and normalize data by the appropriate negative control samples as previously described (Li *et al*., 2020).

A protein summary was first generated where each TMT condition was calculated as a log ratio to the median intensity of all the channels, enabling all channels to have the same denominator. Following calculation of the log_2_ ratio, all samples were normalized by subtracting the median log_2_ ratio (median centering). For each protein, a linear model was used to calculate the following ratio and the corresponding *p*-value:

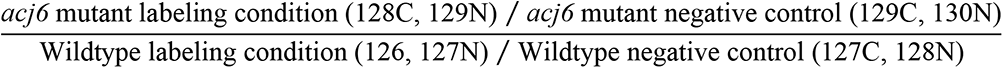

Using log2 transformed TMT ratios, the linear model is as follows:

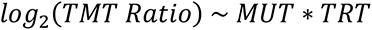

where *MUT* (*acj6* mutant) and *TRT* (treatment) are indicator variables representing *acj6* mutant (*MUT* = 1 for mutant, 0 for wildtype) and labeling condition (*TRT* = 1 for labeled, 0 for negative control) respectively. The above linear model with interaction terms expands to:

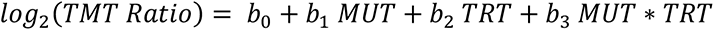

Coefficient *b*_3_ represents the required (log-transformed) ratio between mutant and wildtype conditions taking into account the appropriate negative controls and replicates. A moderated t-test was used to test the null hypothesis of *b*_3_ = 0 and calculate a nominal *p*-value for each protein. These nominal *p*-values were then corrected for multiple testing using the Benjamini-Hochberg FDR (BH-FDR) method (Benjamini and Hochberg, 1995). Figure 2G and Figure S2F show the value of *b*_3_ along the x-axis and the −log_2_(*p*-value) along the y-axis of the volcano plot.

The linear model along with the associated moderated t-test and BH-FDR correction were implemented using the limma library (Ritchie *et al*., 2015) in R.

We note that the ratio compression effect of the TMT strategy reported previously (Savitski et al., 2013; Ting et al., 2011) can also compromise the accuracy of our data, and it is not possible for us to estimate the amount of compression without spiked-in standards. However, by using a less complex sample (proximity-labeled rather than whole cell proteomes) and performing offline fractionation prior to MS analysis, we have reduced ratio compression to the best of our ability without sacrificing the number of proteins identified.

### Quantification of Otk and Piezo expression in PNs

In ImageJ, individual glomeruli were identified based on the NCad staining, and the average FLAG fluorescence intensities for these glomeruli were measured. The intensities were then normalized to the intensity of the *GH146-FLP*-negative DA4m glomerulus, and the normalized values were plotted in Figure 4D and Figure S6B. t-test of independence was used to compare expression of Otk or Piezo between wild-type and *acj6* mutant glomeruli, and *p-*values were adjusted using Bonferroni correction.

### Quantification of adPN dendrite innervation patterns in MARCM-based mosaic analysis

All images of MARCM clones were given human-unidentifiable names and mixed with all other genotypes before scoring. Individual glomeruli were identified using the NCad staining, and then categorical innervation scores were assigned to identified glomeruli. After scoring, the data were imported into python, and the genotype information was revealed.

To test whether overexpressing a transgene rescued any dendrite mistargeting phenotypes caused by the loss of Acj6, we first performed Chi-squared test comparing each glomerulus of a given genotype with that glomerulus in wild-type: if the *p-*value was greater than 0.05, we considered it as fully rescued; if the *p*-value was less than 0.05, we further compared the innervation frequencies of this glomerulus of this genotype with that of *acj6* mutant to see if there was significant partial rescue (*p-*value < 0.05 compared to *acj6* mutant). Note that we did not adjust for multiple comparisons here, because the adjustment would render most mistargeting phenotypes in *acj6* mutant insignificant compared to control and would also fail to detect cases with obvious phenotypic rescue.

---

**Figure S1.**
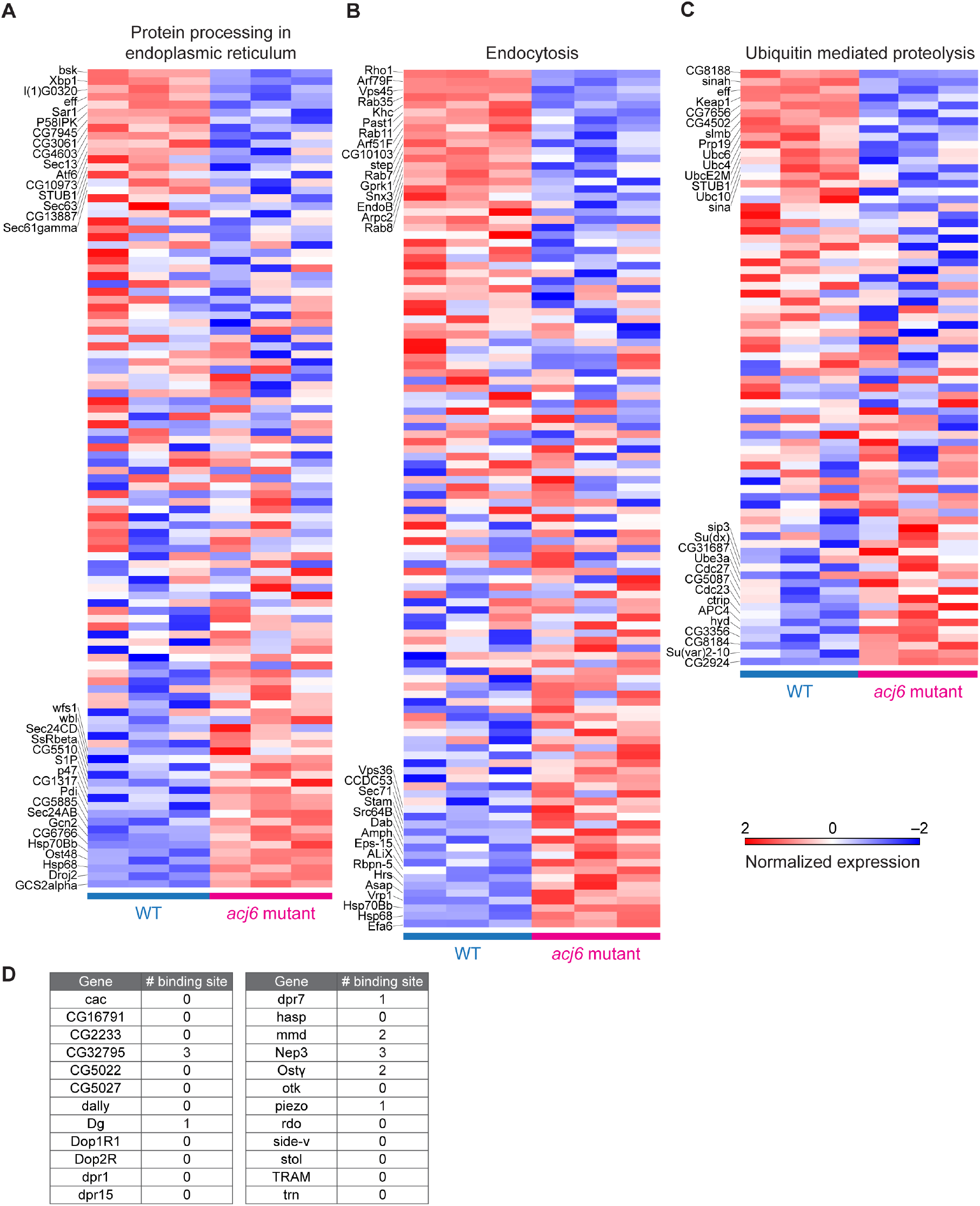
Evidence that Acj6 shapes the PN surface proteomic landscape largely through indirect mechanisms, related to Figures 1 and 2. (A–C) Heatmap of mRNA expression levels from adPN transcriptomes showing the expression of genes with KEGG classification: protein processing in endoplasmic reticulum (A), endocytosis (B), or ubiquitin mediated proteolysis (C; ranked #12 in KEGG classification; *p* = 0.023) in three biological replicates of wild-type (left) and *acj6* mutant (right) samples. Expression is normalized with a mean of 0 and variance of 1 for each gene. (D) Numbers of predicted Acj6 binding sites (Bai *et al*., 2009) found in the promoter region of genes whose PN surface protein expression is regulated by Acj6 (Figure 2G). Note that most predicted binding sites will be non-functional as we identified around 38.5% of all genes in the *Drosophila* genome have predicted Acj6 binding site(s) in their promoter region (futility theorem; Wasserman and Sandelin, 2004), and that even direct transcriptional targets of Acj6 could be further regulated by post-transcriptional mechanisms controlled by Acj6.

**Figure S2.**
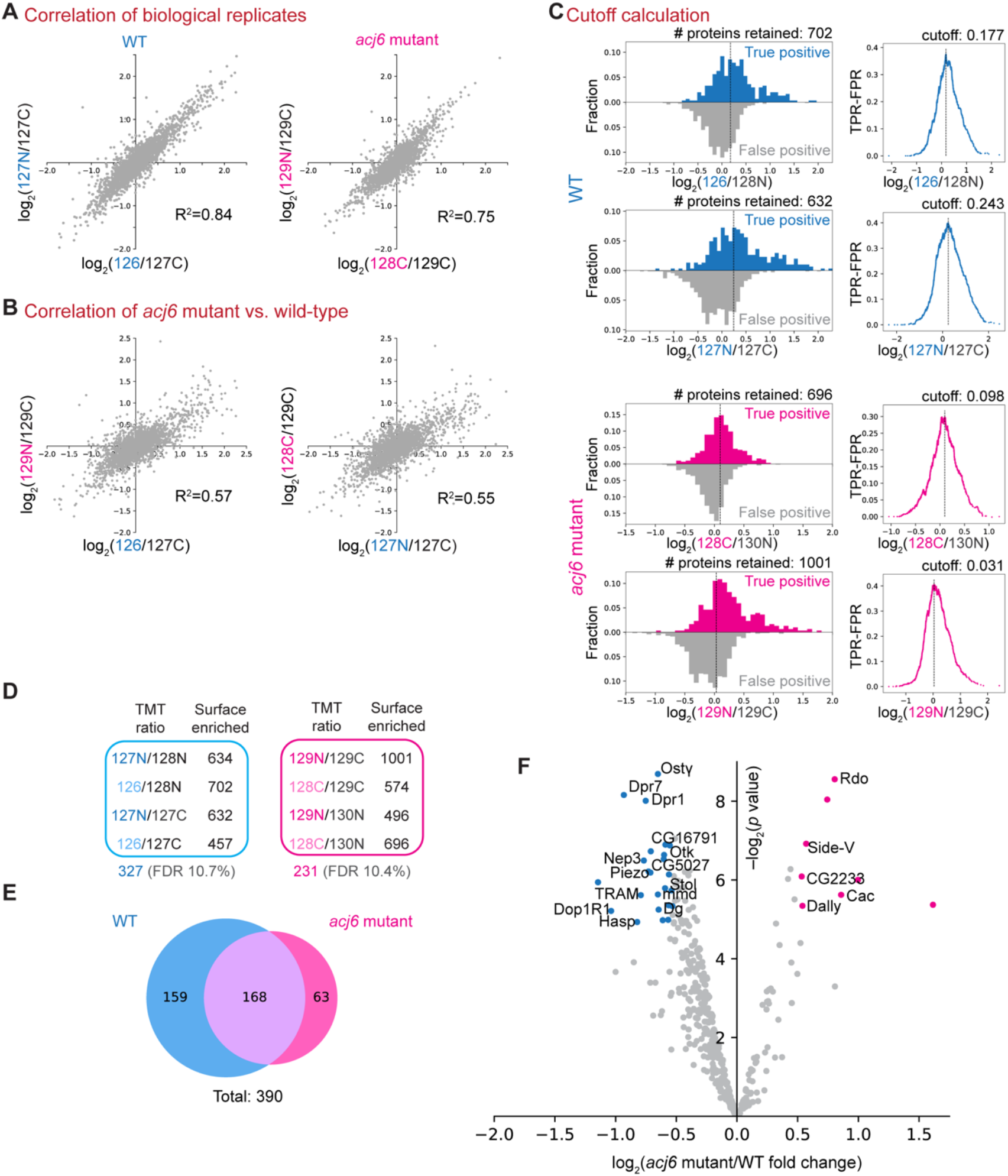
Analysis of PN surface proteomes, related to. **Figure 2** (A) Correlation of biological replicates. (B) Correlation between *acj6* mutant and wild-type samples. (C) Determination of the TMT ratio cutoffs for biological replicates in control and *acj6* mutant. Left: histograms showing the distributions of true positives and false positives by their biotinylation extent. Right: Calculation of cutoff by finding the maxima of [true-positive rate – false-positive rate (TPR – FPR)]. Vertical dashed lines show the cutoff used for each sample. (D) Summary of a more stringent cutoff criterion: a protein must have higher experiment-to-control TMT ratios than the cutoff thresholds in all four possible ratiometric combinations to be included in the final proteome. By this criterion, 327 and 231 proteins were retained in the wild-type (blue) and *acj6* mutant (pink) samples, respectively. (E) Venn diagram showing the size of and overlap between wild-type and *acj6* mutant PN surface proteomes using the more stringent cutoff criterion in (D). (F) Volcano plot showing proteins with altered protein levels on *acj6* mutant PN surface using the cutoff criterion in (D).

**Figure S3.**
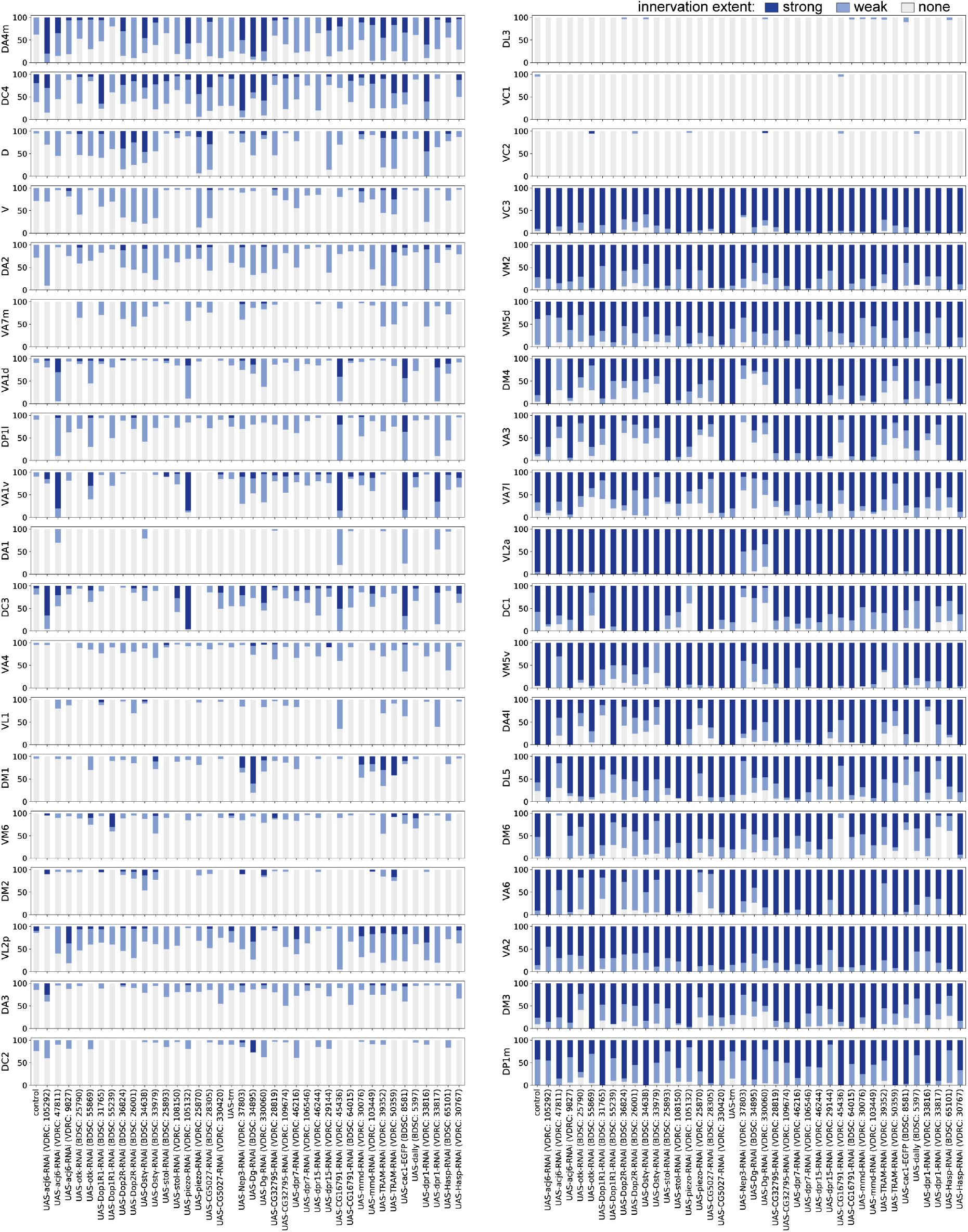
Dendrite innervation patterns of adPNs in the genetic screen, related to. Figure 3 Stacked bar plots summarizing adPN dendrite innervation pattern to each glomerulus (y-axis) when a given candidate gene was overexpressed or knocked down (x-axis). Categorical scorings (top right) were performed blind to genotypes.

**Figure S4.**
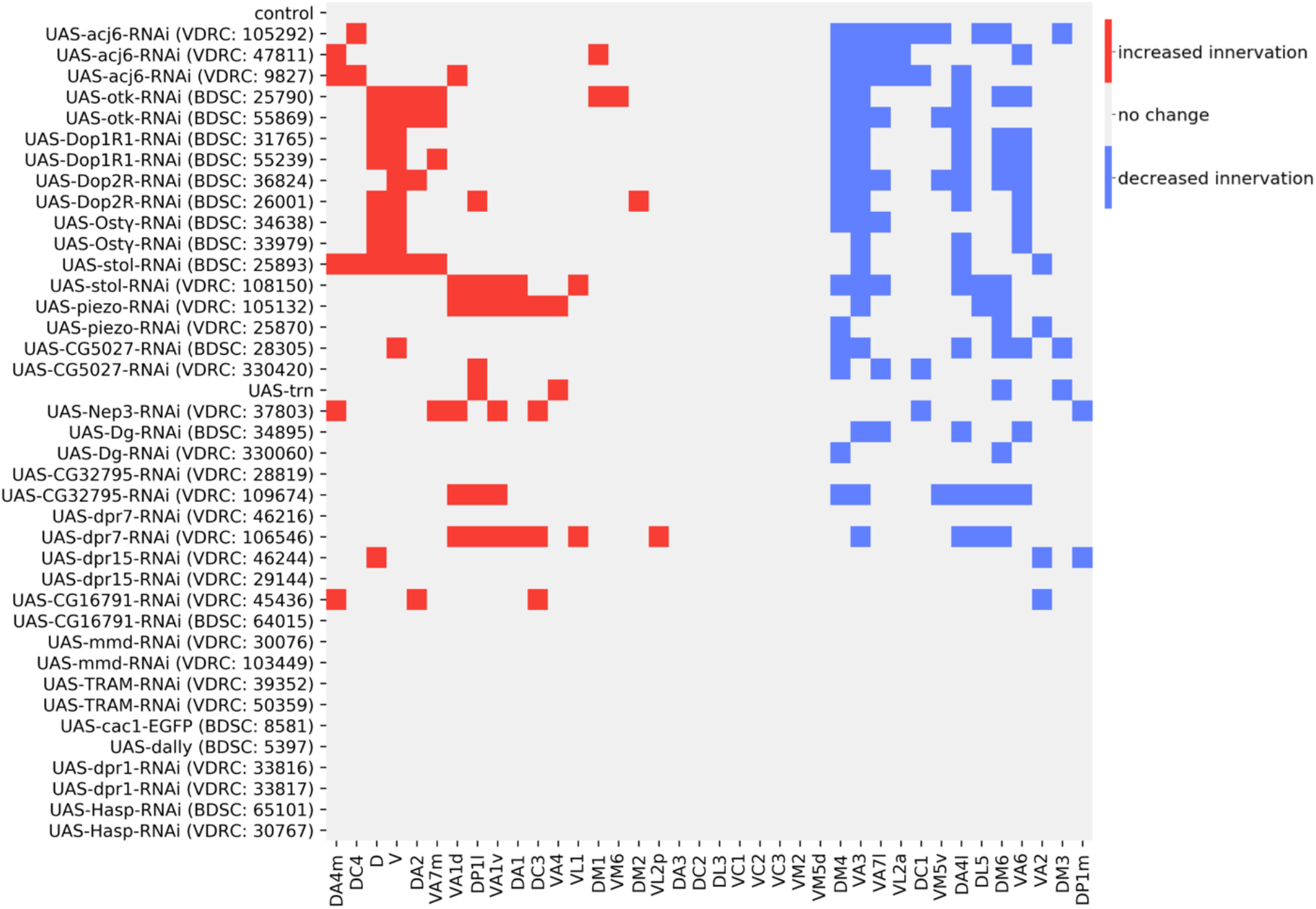
Wiring executors of Acj6 identified from the genetic screen, related to. Figure 3 Heatmap summarizing the Chi-squared test results comparing the innervation extent to each glomerulus in each genotype to that in control (data from Figures S3). Red, ectopic innervation, *p*< 0.05 (adjusted for multiple comparisons). Blue, loss of innervation, *p* < 0.05 (adjusted for multiple comparisons). Note that RNAi-mediated knockdown experiments could suffer from variation in knockdown efficiency and off-target effects, which likely contributed to some phenotypic inconsistency across different RNAi lines targeting the same candidate.

**Figure S5.**
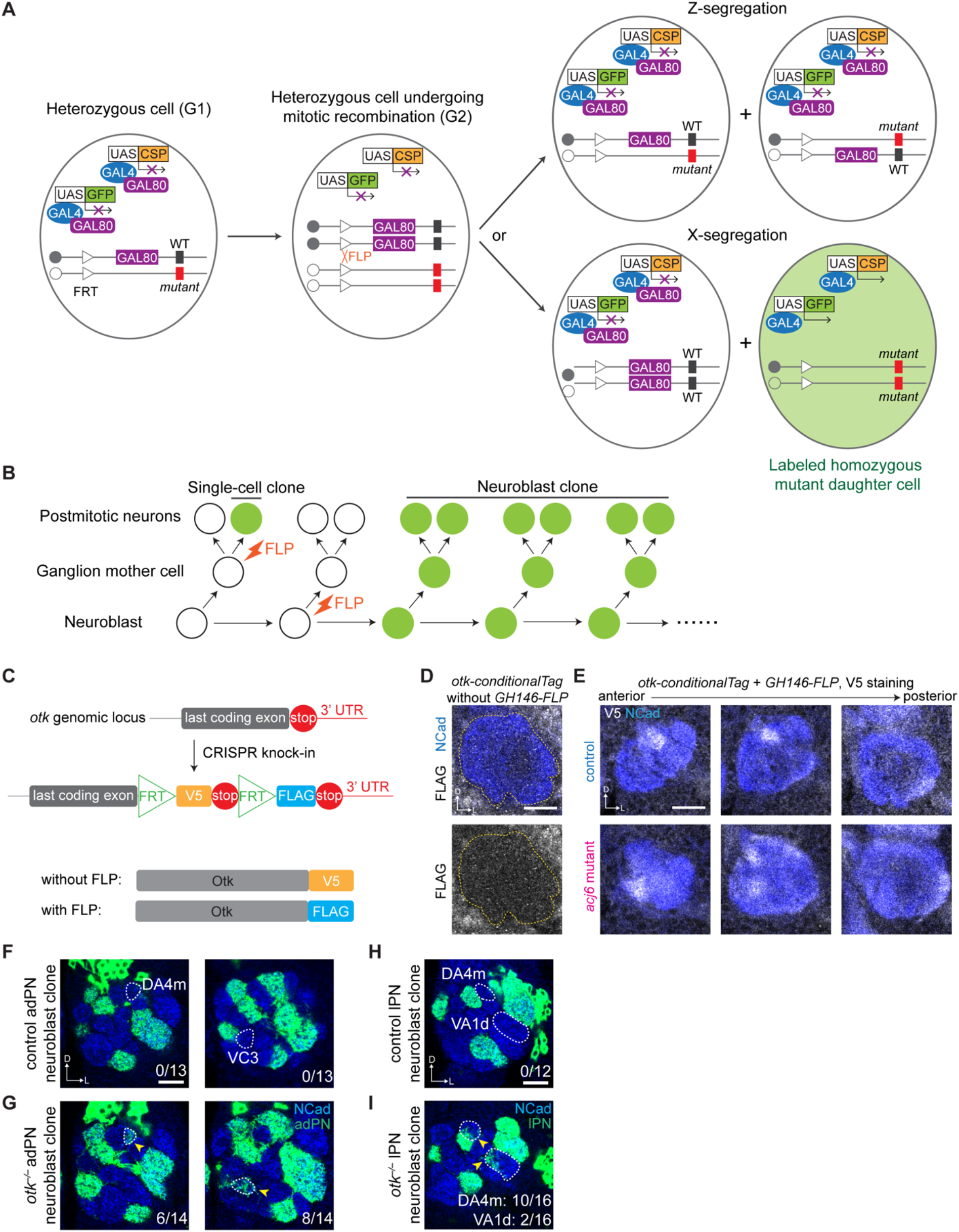
Mosaic analysis scheme, Otk expression pattern, and *otk* mutant phenotypes in neuroblast clones, related to Figures 4–7. (A) Schematic of MARCM analysis. A mutant (*acj6, otk, or piezo*) allele is placed on the same chromosome arm in *trans* to a *GAL80* transgene. Heterozygous cells express GAL80 which represses the activity of GAL4 and is thus unlabeled by GFP. After FLP-mediated mitotic recombination followed by X-segregation (bottom row), one of the daughter cells will be homozygous for the mutant and will lose *GAL80*. Those homozygous mutant cells will be labeled with GFP and can also express one or two candidate cell surface proteins (CSP) for the rescue assay. Mitotic recombination followed by Z-segregation (top row) does not generate daughter cells with altered genotype or transgene expression. (B) Illustration of cell division patterns in a neuroblast lineage. MARCM can be used to generate GFP-labeled PN single-cell or neuroblast clones. All clones were induced by heat shock applied in newly hatched larvae (0–24 hr after larval hatching), so our analyses are restricted to the larval-born adPNs, lPNs, and DL1 single-cell clones (Jefferis *et al*., 2001). (C) Endogenous conditional tagging of Otk to reveal its cell-type-specific protein expression pattern. (D) *otk-conditionalTag* has minimal FLAG background in the antennal lobe (outlined in yellow) without FLP expression. (E) V5 staining showing the expression of Otk in the antennal lobe contributed by *GH146-FLP*- negative cell types, including ORN axons, local interneurons, glia, and a small fraction of PNs that are not covered by *GH146-FLP*, such as DA4m PNs. (F and G) DA4m and VC3 glomeruli were ectopically innervated by *otk^−/−^* adPN neuroblast clones (G) compared to wild-type (F). (H and I). DA4m and VA1d glomeruli were ectopically innervated by *otk^−/−^* lPN neuroblast clones (I) compared to wild-type (H). Scale bars, 20 μm. D, dorsal; L, lateral. The number of clones with mistargeting phenotype over the total number of clones examined is noted at the bottom right corner of each panel.

**Figure S6.**
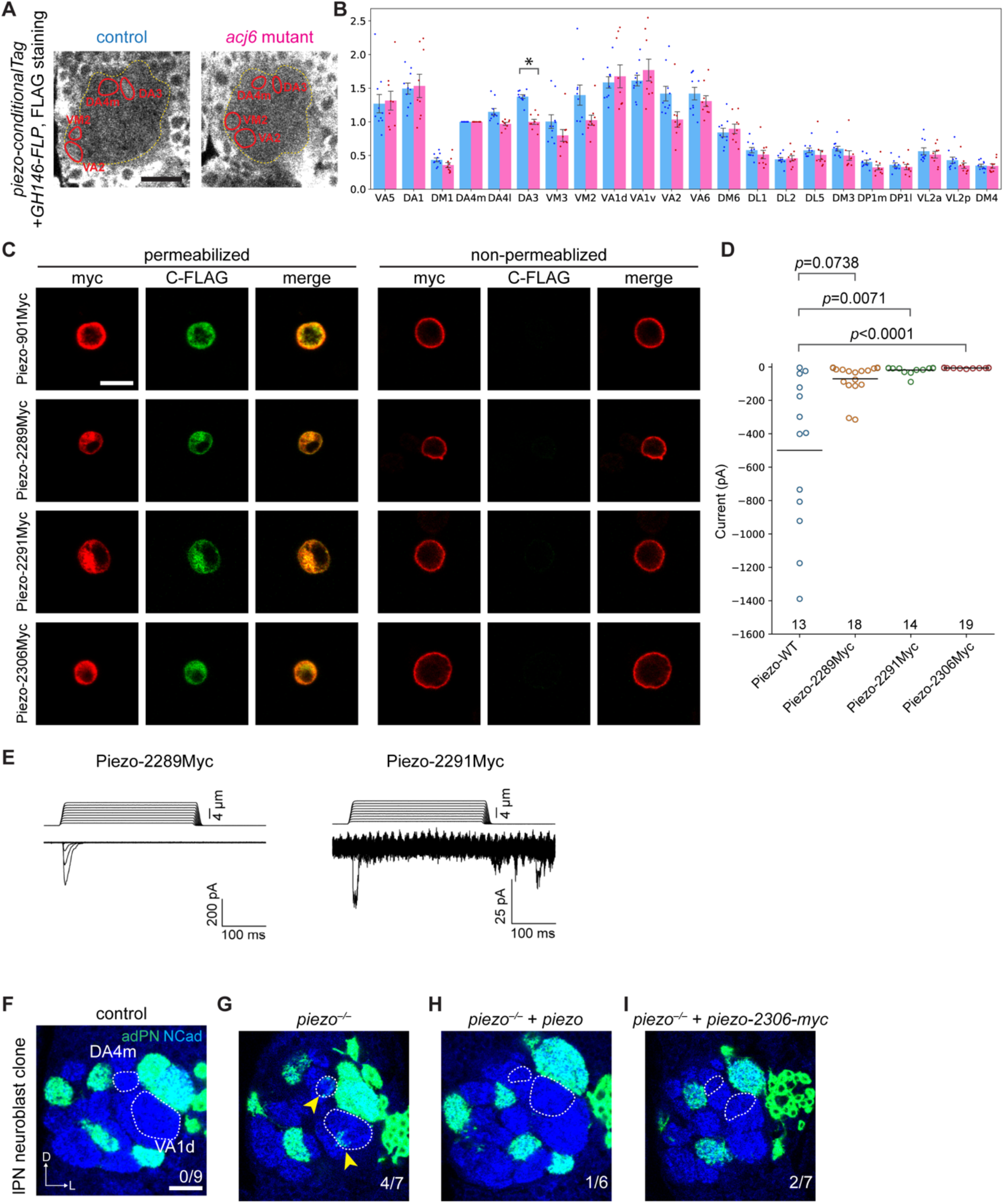
Piezo expression pattern, characterization of myc insertion mutants, and *piezo* mutant phenotypes in lPN neuroblast clones, related to. **Figure 5** (A) Expression of Piezo in PNs at 42–48 hr APF in wild-type or *acj6* mutant brains. Antennal lobe (yellow) and the DA4m, DA3, VM2, and VA2 glomeruli are outlined. *GH146-FLP* was used to express FLP in the majority of PNs. (B) Quantification of Piezo expression in developing PNs of wild-type (n = 10) and *acj6-*mutant (n = 9) animals. Fluorescence intensities were normalized to the DA4m glomerulus (*GH146-FLP* negative) in each antennal lobe. Mean ± s.e.m. *: *p* < 0.05 (two-tailed t-test; *p*-values were adjusted for multiple comparisons). (C) Piezo myc insertion mutants have normal cell-surface expression in *Drosophila* S2 cells. Representative images of non-permeabilized and permeabilized staining of the extracellular myc tag (red) and the intercellular C-terminal FLAG tag (green). dPiezo-901Myc is a positive control to show wild-type Piezo expression. Scale bar, 10 μm. (D) Scatter plot represents mechanically activated whole-cell I_max_ currents recorded at –80mV from human *PIEZO1* knockout HEK 293T cells transfected with wild-type *piezo* or three *piezo* Myc insertion mutants. The number of cells recorded, mean, and *p*-values of Dunn’s multiple comparison test (compared to wild-type) for each construct are shown. (E) Representative stimulus and current traces of indentation-induced whole-cell currents from *piezo-2289Myc* or *piezo-2291Myc* transfected cells are shown. (F–I) Dendrite innervation patterns of control (F), *piezo^−/−^* (G), *piezo^−/−^, UAS-piezo* (H), and *piezo^−/−^, UAS-piezo-2306Myc* (I) lPN neuroblast clones. Scale bar, 20 μm. D, dorsal; L, lateral. The number of clones with mistargeting phenotype over the total number of clones examined is noted at the bottom right corner of each panel.

**Figure S7.**
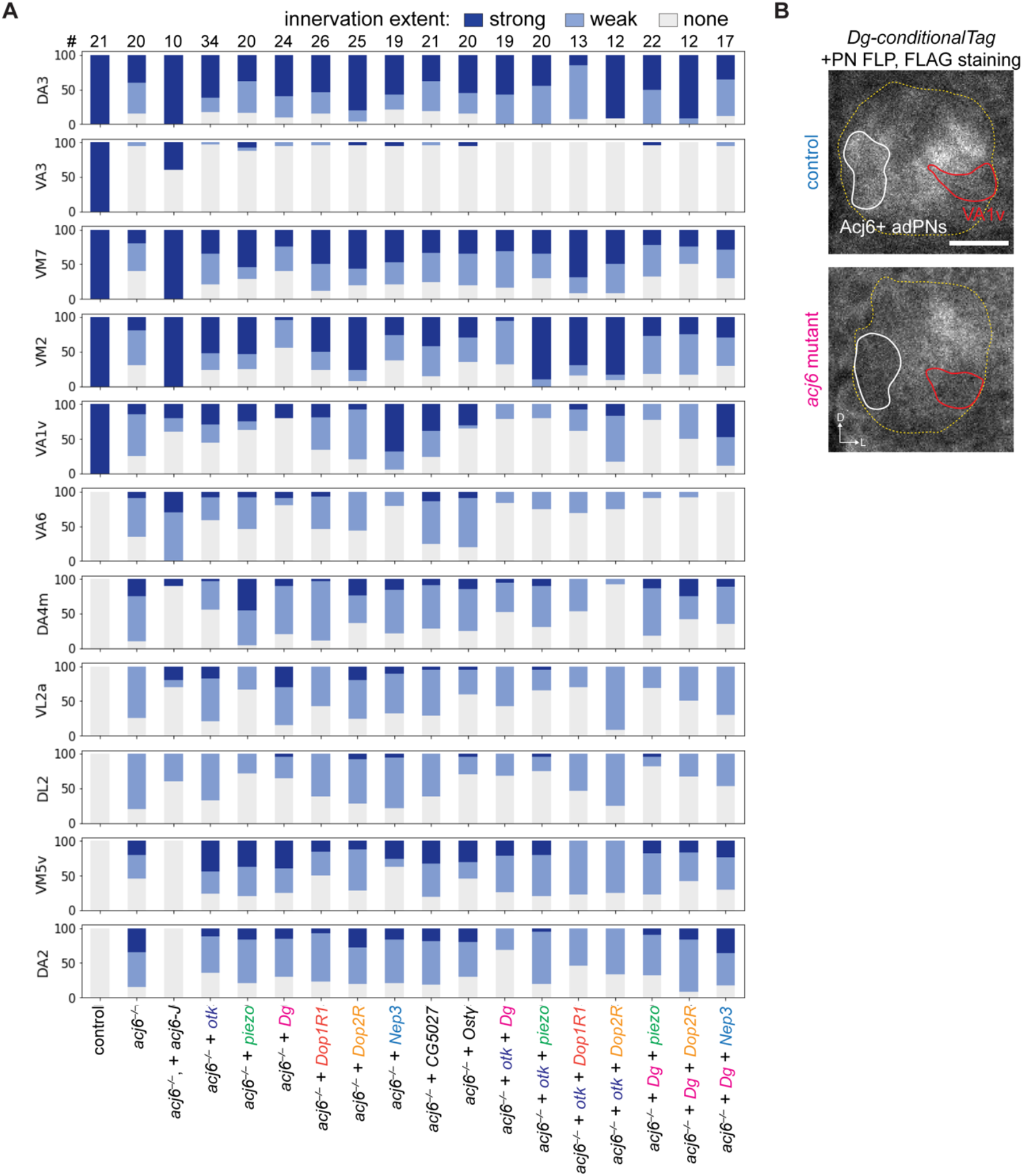
Dendrite innervation patterns of adPNs in the epistasis experiments, related to. **Figure 6** **and** **Figure 7** (A) Stacked bar plots summarizing dendrite innervation pattern of adPN neuroblast clones to each glomerulus in each genotype. Confocal stacks of different genotypes were scrambled and blinded during scoring. Total number of adPN neuroblast clones examined for each genotype is labeled on top. (B) Dg conditional tag staining shows that Dg is not expressed in VA1v PNs (red outline) at 42– 48 hr APF. Many medial glomeruli innervated by adPN dendrites (white outline) have decreased Dg level in *acj6* mutant animals. Scale bars, 20 μm. D, dorsal; L, lateral. *GH146-FLP* was used to express FLP in the majority of PNs.

## SUPPLEMENTAL TABLES

**Table S1. Processed RNA-seq data rank ordered according to adjusted P-value of the comparisons between *acj6* mutant and wild-type samples (see separate Excel sheet)**

**Table S2. Raw and processed proteome data** (see separate Excel sheet)

**Table S3.**
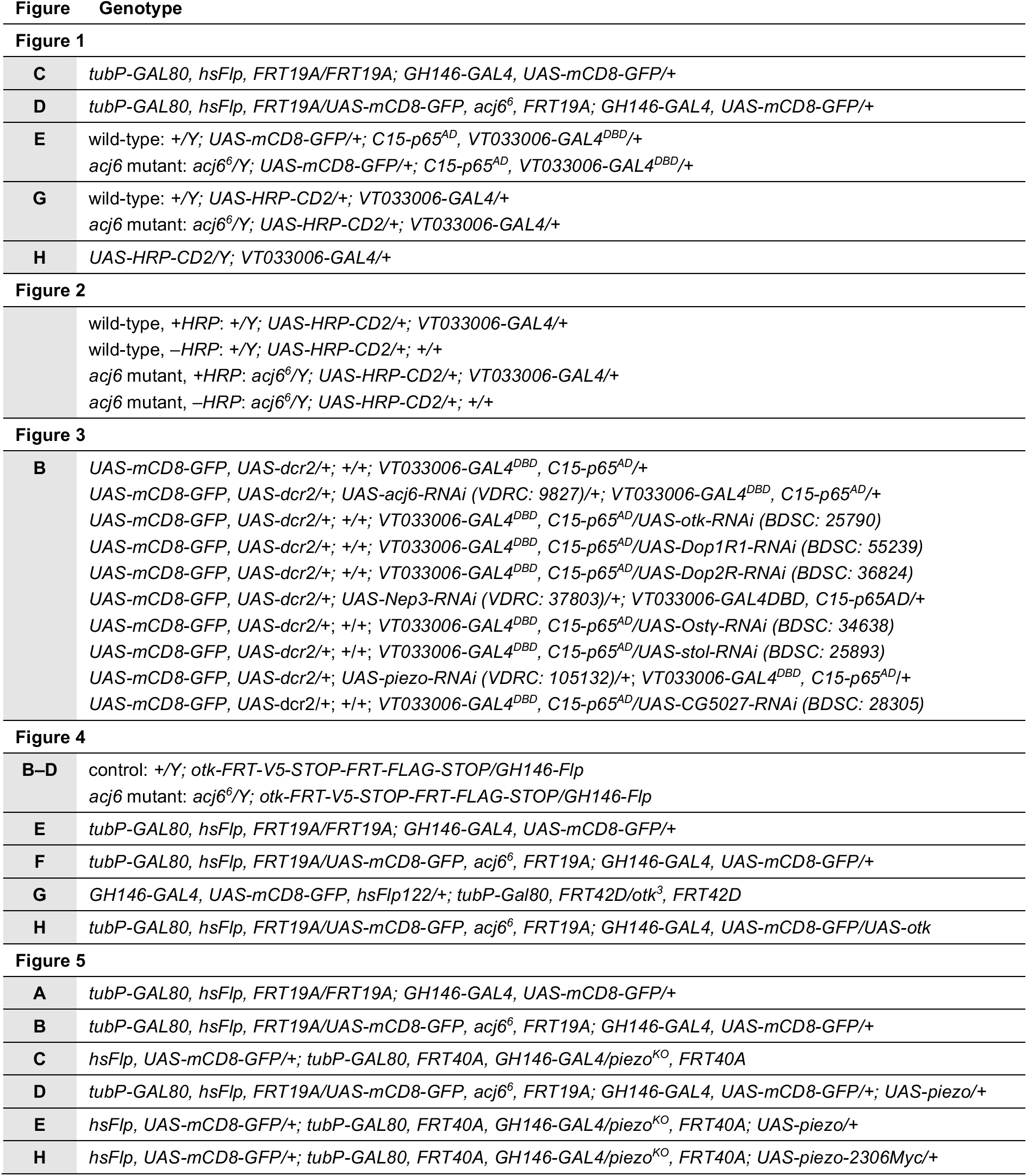
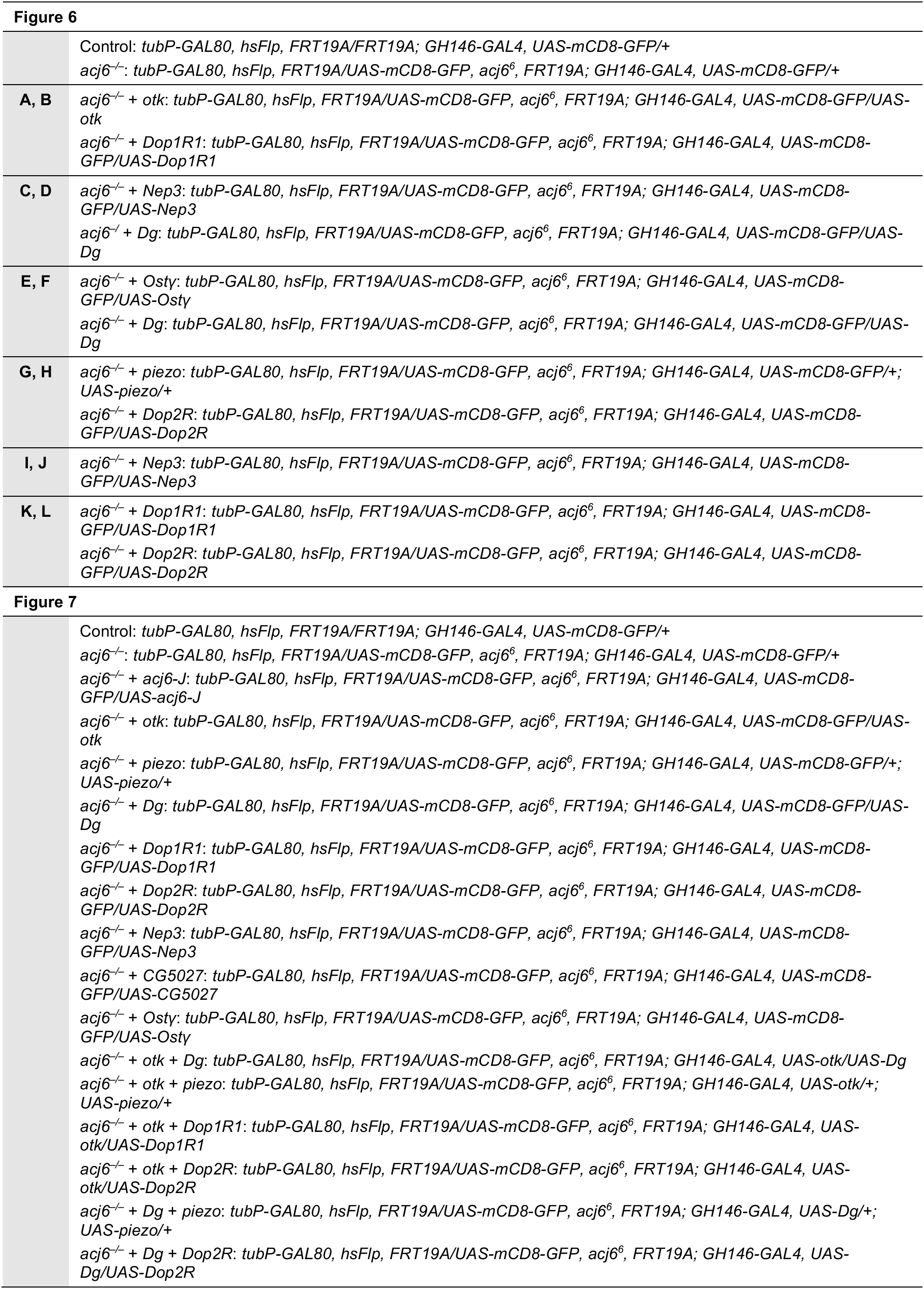
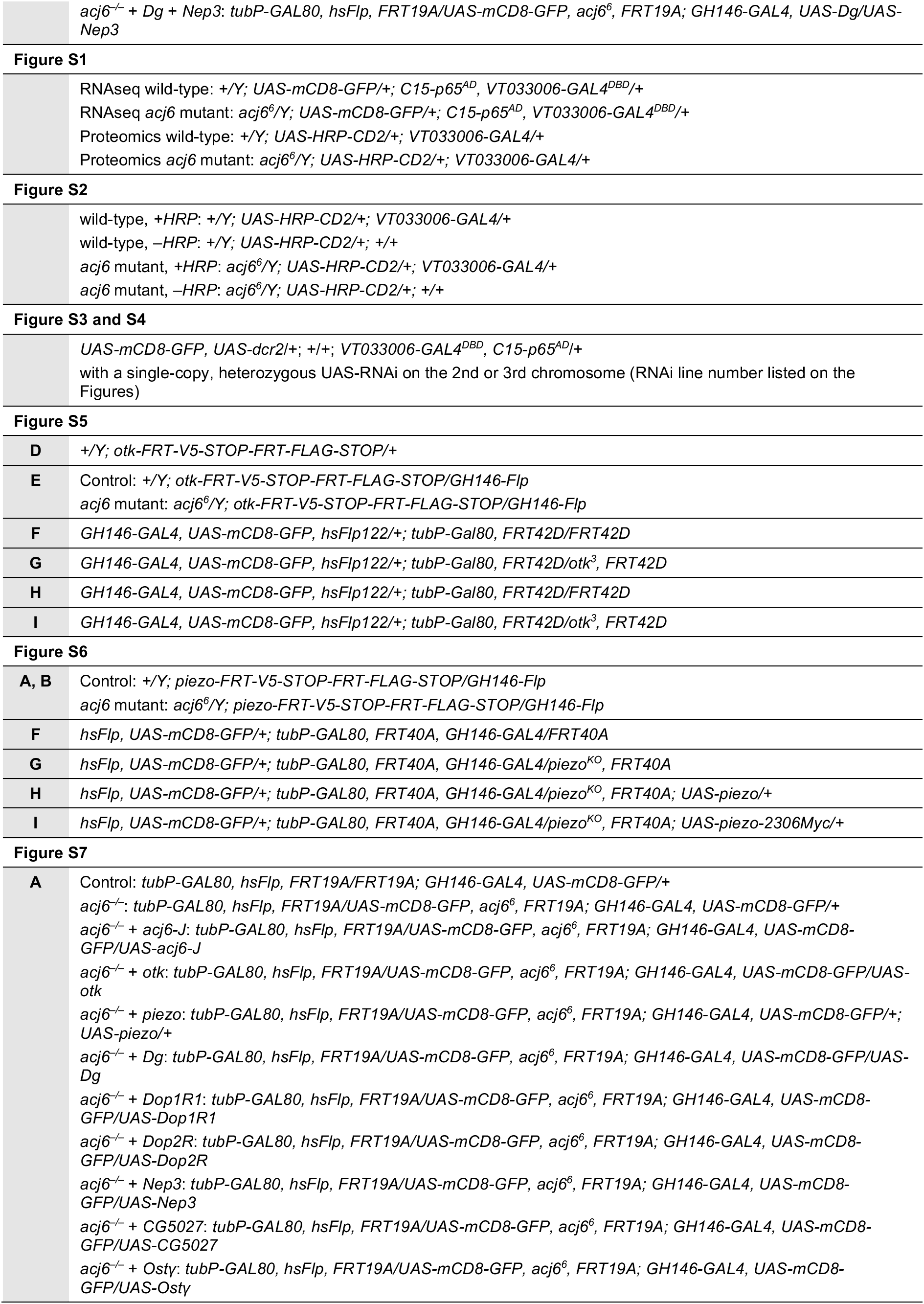
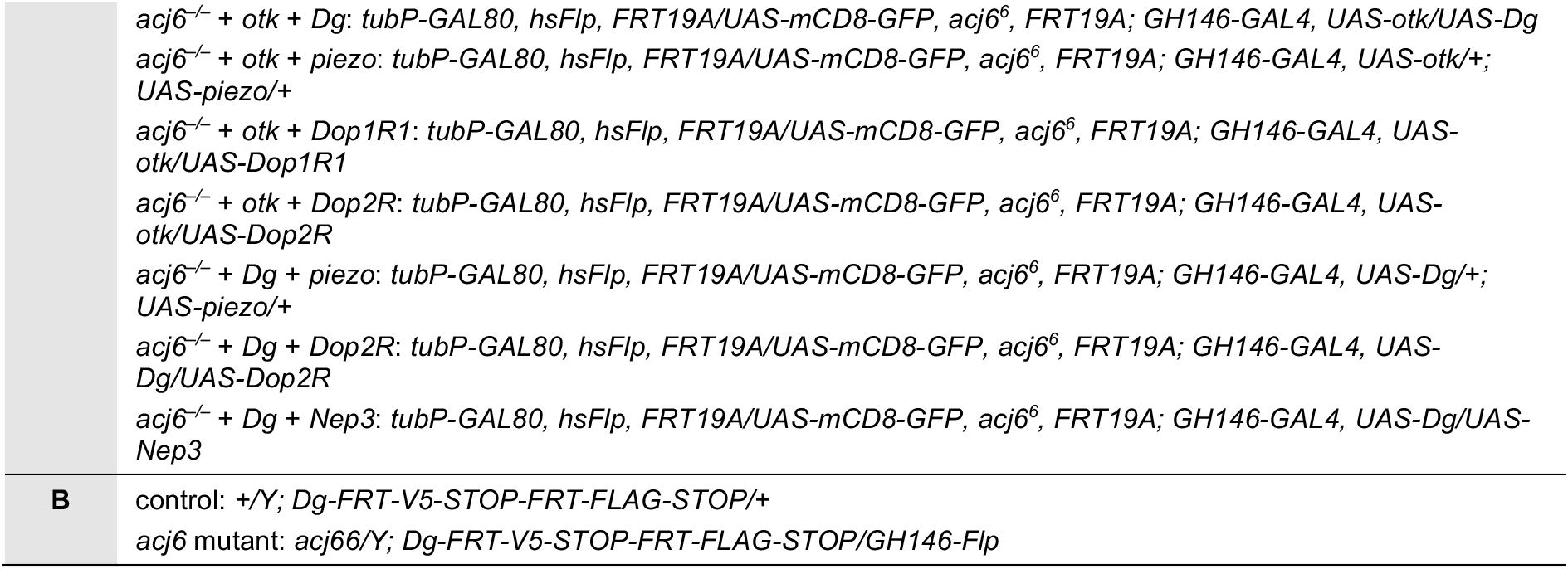
Genotypes of flies in each experiment.

